# Barley miRNAs and their targets regulation in response to heat stress at the early stage of development

**DOI:** 10.1101/2024.10.25.620191

**Authors:** Katarzyna Kruszka, Andrzej Pacak, Aleksandra Swida-Barteczka, Jacek Kesy, Artur Jarmolowski, Zofia Szweykowska-Kulinska

## Abstract

MiRNAs are key regulators of gene expression controlling plant development and response to environmental stresses. In this work we studied global dynamics of accumulation of conserved and identified novel barley miRNAs at early stage of plant development during heat stress (1h, 3h and 6h of heat stress). The majority of miRNAs responds to heat stress after 3h and 6h of heat stress duration (124 and 155, respectively). The comparison of heat-induced changes in mature miRNA accumulation to their cognate precursor levels allowed to indicate a smaller group of miRNAs that are controlled at transcriptional level and a larger group that is controlled posttranscriptionally in response to heat stress. For miRNAs with the significant accumulation changes during heat treatment, target mRNAs were identified. Moreover, novel targets have been experimentally assigned for selected miRNAs. mRNA of the effector protein of miRNA activity, AGO1B was found to be downregulated by increased miR168 during heat stress. Importantly, miRNA/mRNA target module miR399c/PHO2 responsible for the phosphorus uptake exhibits dynamic changes under heat stress conditions suggesting adaptation of plant development to stress conditions. This study provides new data for developing miRNA and their mRNA target-based strategies in barley breeding in response to heat stress.

**Keypoints:**

- Involvement of small RNAs in response to the heat stress conditions have been studied in young barley plants.
- The largest number of heat responsive miRNAs was found after 3h and 6h of heat duration.
- Heat-induced mature miRNA accumulation and their precursor levels showed complex transcriptional or posttranscriptional regulation.
- MiRNA-target modules responsive to heat stress were identified

## Introduction

MicroRNAs (miRNAs) are small, single stranded RNA molecules, mostly of 20-22 nt long [1]. MiRNAs are the most prominent part of posttranscriptional gene expression silencing (PTGS) mechanism. MiRNAs designate target mRNAs to cleavage or translational inhibition, depending on the level of complementarity between miRNA and target mRNA [2, 3]. Therefore, generally the biological role of miRNAs is to downregulate gene expression. MiRNAs control many aspects of plant development, like the biomass growth, leaves and roots growth, induction of flowering or flower patterning [4–7]. Also, plant adaptation to many abiotic and biotic stresses is driven by miRNAs and their target mRNAs [8–10].

Plant miRNAs usually originate from individual miRNA genes (*MIRs*) and are transcribed by RNA polymerase II (RNA Pol II) [11, 12]. Therefore the primary miRNA transcripts (pri-miRNAs) are 5’ capped and 3’ polyadenylated. A specific exception from this rule is a group of *Cuscuta campestris* miRNAs called interface-induced miRNAs, transcribed by RNA Pol III [13]. Characteristic feature of the pri-miRNA is the presence of a hairpin structure (pre-miRNA) in which miRNA and miRNA* are imbedded. DICER-LIKE1 (DCL1) catalyzes two sequential, endonucleolytic cleavages of the pri-miRNA releasing finally the miRNA in a duplex with miRNA* [14, 15]. There are two ways by which DCL1 can approach pri-miRNA. In the base-to-loop mechanism, DCL1 performs the first cleavages at the base of the pre-miRNA stem, releasing it to the nucleoplasm [16]. In the second mechanism pri-miRNA is processed by the DCL1 at the loop-to-base direction. SERRATE (SE) and HYPONASTIC LEAVES 1 (HYL1) ensure the cleavage efficiency and precision and together with DCL1 form the core of microprocessor complex [17]. 3’ ends of the miRNA/miRNA* duplex are protected from degradation by HUA ENHANCER1 (HEN1)-driven 2’-*O*-methylation [18]. The miRNA/miRNA* duplex is loaded into ARGONAUTE1 (AGO1) and exported from nucleus to cytoplasm by CHROMOSOMAL MAINTENANCE1/EXPORTIN1 (CRM1/EXPO1) [19]. The miRNA* is then removed and miRNA together with AGO1 forms RNA-induced silencing complex (miRISC) capable to cleave target mRNA [20].

The efficiency of miRNA biogenesis is regulated at multiple levels. *MIR* expression is regulated by transcription factors that may stimulate or repress transcription of *MIR* genes [11, 21, 22]. An interesting mode of *MIR* expression enhancement is the R-loop formation near the transcription start site region [23]. Many *MIR* genes possess introns. Intron presence and its splicing is necessary for effective mature miRNAs accumulation like in the case of exonic miRNA163 and intronic miRNA400 and miRNA402 [24–27]. Intron removal is mandatory for folding of pre-miRNA of monocotyledonous specific miRNA444 family members [28, 29]. Another distinctive factor influencing miRNA maturation is miRNA/miRNA* duplex formation within the pre-miRNA stem. It strongly depends on the presence of N^6^-methylation of adenosine (m6A) in the duplex and consequently the lack of m6A leads to decreased level of selected mature miRNAs [30].

Barley, *Hordeum vulgare*, is the fourth most cultivated cereal crop worldwide [31]. Heat stress negatively influences global crop production including barley which is considered to be one of the less heat stress sensitive cereals [32]. However, detailed studies confirmed that the generative phase of barley development is severely affected by heat, especially, floret development is particularly vulnerable to aberrations. Elevated temperatures mainly cause abnormal anthers and ovules formation. The overall effect is heat stress induced sterility [33]. High temperatures negatively influence the duration of grain filling what results in reduced overall grain weight. Also, the barley grain quality is substantially lowered by altered specific traits like starch, β-glucan and nitrogen content [34]. So far, studies concerning negative heat stress influence on barley were conducted mainly during reproductive growth stage. There are almost no data concerning heat stress induced changes in barley early development.

For barley, only four heat-responsive miRNAs were reported [35]. In the case of other cereals, the most comprehensive studies of miRNAs accumulation during heat were conducted for wheat. In two wheat cultivars, heat tolerant, RAJ3765, and heat susceptible, HUW510, 84 conserved and 93 novel miRNAs were identified, where 32 and 40 conserved and 15 and 18 novel miRNAs were differently accumulated in RAJ3765 and HUW510 cultivars, respectively [36]. In wheat, cv. Chinese spring, 104 miRNAs was identified and 36 of them were differently accumulated [37]. Both these studies were conducted in booting stage of plant development. Heat stress applied at night during booting of rice resulted in 102 differently accumulated miRNAs between heat tolerant and heat susceptible rice cultivars. Forty nine of these miRNAs are suggested to have significant influence on the difference between the heat tolerance level [38]. Studies conducted on maize tissues collected in three-leaf and the beginning of flowering developmental stages revealed 18 miRNAs and 15 miRNA* species differently accumulated during heat stress [39]. Importantly, it was shown that simple sequence repeats (SSRs) found in *MIRs* can be used as molecular markers differentiating between heat tolerant and heat susceptible wheat cultivars. Therefore, miRNA-derived SSRs are a new tool for marker assisted heat-tolerant cereals breeding programs [40].

The goal of this study was to reveal the involvement of miRNAs and their targets in 2-week-old barley plants in response to heat stress. Our findings open new perspectives for developing new strategies using miRNAs and their targets in barley breeding in response to heat stress.

## Material and methods

### Plant material and growth conditions

Spring barley plants, cultivar Rolap seeds were received from the Institute of Plant Genetics of the Polish Academy of Sciences (Poznan, Poland) [41]. Plants were grown in a Conviron environmental chamber, BDW120 High-Light Plant Growth Room (Conviron, Winnipeg, Manitoba, Canada) with a 16 h day/8 h night photoperiod, and 600 μmol · m^-2^ · s^-1^ light conditions. Plants were grown at 21 °C day/15 °C night temperature in 250 ml pots containing field soil, and were watered to maintain optimal growth conditions of 70% SWC (Soil Water Content). Two-week-old plants at the three-leaf stage (code13 of Zadoks system; [42]) were subjected to heat treatment at 35.5 °C. Control plants were grown at a constant temperature of 21 °C. Plants were collected after 1, 3 and 6h of heat stress in three biological replicates.

### RNA isolation and small RNA library preparation

A total RNA was isolated from 100 mg of the ground tissue using TRIZOL reagent [43] accompanied by Plant RNA Isolation Aid (Thermo Fisher, Vilnius, Lithuania). Mixture was purified using Direct-zol RNA Miniprep kit (ZymoResearch, Irvine, CA, USA). The quality of the isolated RNA was verified using an Agilent 2100 Bioanalyzer and Nano Plant RNA assay (Agilent, Vilnius, Lithuania). For each time point and condition independent small RNA (sRNAs) libraries were prepared. Small RNAs of 15–30 nt in length were separated on denaturing 8 M urea 15% polyacrylamide gel, purified and ligated to 3′ and 5′ RNA adapters derived from TruSeq Small RNA Library Preparation Kit (Illumina, San Diego, CA, USA). Small RNA libraries were created using TruSeq Small RNA Library Preparation Kit (Illumina, San Diego, CA, USA). Quality and quantity of the each library was analyzed using an Agilent 2100 Bioanalyzer and Nano Plant RNA assay (Agilent, Vilnius, Lithuania) for quality control, and Qubit 3.0 Fluorometer measurement for estimation of each library concentration. Eighteen libraries were sequenced (nine libraries were pooled and put on one lane). Sequencing (2×50+8) was performed using NovaSeq 6000 machine (Illumina) by Fasteris S. A. (Switzerland).

### Quantitative Real-Time RT-PCR (RT-qPCR)

3 μg of DNA-depleted RNA were reverse-transcribed with Invitrogen SuperScript III Reverse Transcriptase (ThermoFisher Scientific, Waltham, MA, USA) and 0.5 μg Oligo(dT)_18_ Primer (ThermoFisher Scientific, Waltham, MA, USA). cDNA samples were 4-fold diluted and 1 μl was used as template. Quantitative Real-time PCR was performed with Power SYBR Green PCR Master MIX (Applied Biosystems, Warrington, UK) and two specific primers (final concentration 500 nM each) using a QuantStudio 7 Flex Real-Time PCR System (Applied Biosystems) in 10 μl of reaction volume, in 384-well plates. Each real-time PCR reaction was performed independently for three biological replicates. The barley ADP-ribosylation factor 1-like (GenBank: AJ508228.2) transcript fragment was simultaneously amplified and detected as an internal reference [44]. Expression levels were calculated with the relative quantification method (2^-ΔΔCt^) as a fold-change value. The significance of the changes was tested with a Student’s t-test. Primer sequences can be found in Table S1.

### Degradome preparation and analysis

Degradome library preparation was performed according to the previously described approaches [45, 46]. Degradome libraries were sequenced by Fasteris S. A. (Switzerland). Degradome analysis and T-plots construction was performed using PAREsnip2 software [47].

### Auxin level measurement

Control and heat-stressed plants for IAA (indole-3-acidic acid) determination were used from the heat experiment described above. 100mg of tissue crushed in liquid nitrogen was incubated overnight in a cold room in a mixture of 80% acetonitrile with 5% formic acid and 1 mM BHT and an internal standard, deuterium-labelled IAA (d2-IAA, 4 ng/sample). Mixture of solid magnesium sulfate and sodium chloride (3:1) in amount of 0.3g was added, and samples were vortexed for 5 min and then centrifuged for 10 min at maximum speed (14,000 rpm). Sodium sulfate was added to the obtained supernatants, and the samples were vortexed and centrifuged as described above.

The obtained supernatant was dried under a stream of nitrogen at 50°C and the remaining residue was suspended in 1 M formic acid. The samples were then extracted using solid-phase C18 octadecyl columns (J.T. Baker C18 columns #7020–01), dried under a stream of nitrogen at 50°C, and the remaining residue was suspended in 100 μl of 80% methanol, followed by suspension in 35% methanol. IAA concentration was determined using ultra-high pressure liquid chromatography with tandem mass spectrometry (UHPLC–MS/MS; Shimadzu Nexera XR UHPLC/LCMS-8045 system; Kyoto, Japan) equipped with an Ascentis Express C-18 column (2.7 μm, 100 × 2.1 mm, Supelco, United States). Then, the obtained peak surfaces of standard and endogenous ABA were exported to an Excel file and analyzed using the formula: (endogenous IAA peak surface/internal standard surface) × 10/weight of used tissue in grams.

### Bioinformatic analyses

Transcript sequences for barley target genes were obtained from the Ensembl Plants database [48]. Small RNA-seq data were analyzed using CLC Genomics Workbench 21.0.5 (QIAGEN Aarhus A/S). Only reads R1 from paired reads were analyzed. Adapters were removed in two steps: (i) by automatic adapter read-through trimming together with low quality sequence removal (limit=0.05) and ambiguous nucleotides removal (max. 2 ambiguous nucleotides), (ii) by adapter sequences trimming together with low quality sequence removal (limit=0.05), ambiguous nucleotides removal (max. 2 ambiguous nucleotides) and removal of sequences on length: minimum length 18 nucleotides and maximum length 25 nucleotides. Counts were normalized per 1 000 000 reads. Shoot samples analyses were performed by two group comparison (samples treated by high temperature vs. control temperature for 1h, 3h and 6h collection sample time). Small RNA sequences were annotated to miRbase Sequence database (release 22.1) without mismatches and with strand-specific alignment. EDGE (Empirical analysis of DGE) method was used to find statistically significant fold changes in small RNA abundance levels between samples derived from different temperature exposed samples. P value correction was carried out using EDGE FDR p-value and EDGE Bonferroni correction calculation. For further analysis miRNAs with length of 20-22 nt, with read score more than 10 and pval ≤ 0.05 were selected. For novel microRNA EDGE analysis was performed using set of 20-22 nt small RNA.

Secondary structures of barley pre-miRNAs were predicted using Folder Version 1.11 software with RNAfold algorithm [49]. Stem-loop structures with the lowest minimal folding free energy (ΔG kcal/mol) were shown in Table S2.

### Deposited data

The small RNA-seq data discussed in this publication have been deposited in NCBI’s Gene Expression Omnibus, GEO [50] and are accessible through GEO Series accession number GSE272607 [51].

## Results

### Identification of heat responsive miRNAs in 2-week-old barley plants subjected to 1h, 3h, and 6h of heat stress

Spring barley plants genotype Rolap were grown for 2 weeks up to three-leaf stage (code 13 of Zadoks system) at constant temperature of 21 °C. Afterwards, half of the plants was subjected to heat treatment at 35.5 °C. Control and stressed plants were collected at 1, 3 and 6h time points in three biological replicates.

To identify conserved and unique, previously published barley miRNAs involved in the response to high temperature stress small RNA libraries were prepared from RNA isolated from control and heat-treated plants (from all three time points of control and heat duration; altogether, 18 libraries). After the sequencing reaction, depending on library analyzed, we obtained between 4 600 - 4 750 miRBase annotated sRNAs (miRBase, release 22.1, [52]. Illumina sRNA sequencing was performed and the analysis of changes in the level of small RNAs 20-22 nt long was carried out. Levels of 65, 124 and 155 conserved miRNAs were changed after 1h, 3h and 6h, respectively (Table 1, Table S3,). In the case of 1h heat stress duration the most pronounced differences in accumulation levels were observed for miRNAs belonging to miR159, 169 and 444 families (Figure 1). After 3h the strongest effects were observed for miRNAs belonging to miR156, 169, 444 families, while 6h of heat caused the most profound changes in the accumulation of miRNAs from miR160, 169 and 444 families. MiR444c* is the most abundant miRNA in 1h and 3h heat stressed plants, while after 6h of stress miR167f was observed at the highest level (Figure 1). Some miRNAs that were found to be affected in their level after 1h of stress were also changed at 3 and/or 6h of heat duration. Thus after 6h of heat we partly observe a cumulative effect of changes that started already after 1h of heat stress (miR166f, miR168 and miR9662) (Table S3).

**Table 1.** Summary of accumulation changes of conserved miRNAs after 1h, 3h and 6h of heat stress treatment.

| miRNA level | 1h | 3h | 6h |
| --- | --- | --- | --- |
| all diferrentially accumulated | **65** | **124** | **155** |
| down regulated | 39 | 80 | 73 |
| up regulated | 26 | 44 | 82 |
| miRNA families | 20 | 26 | 36 |

**Figure 1.**
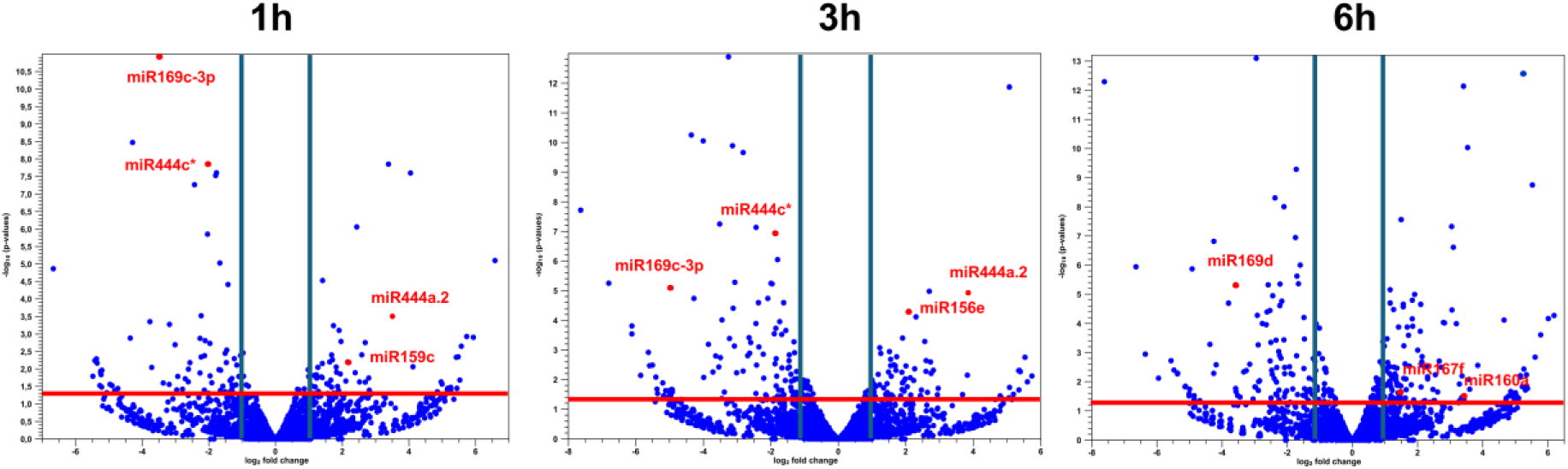
Volcano plots of the differentially expressed barley miRNAs after 1, 3 and 6h of heat stress treatment. miRNAs marked on the plots represent the most changed miRNAs at given time point. The horizontal axis represents the log2 fold change and the vertical axis represents p value (−log10). Vertical lines correspond to 2-fold up-and downregulation, and the horizontal red line represents pval= 0.05. Red dot corresponds to miRNA with maximal 8-fold FC (fold change).

Analysis of mature miRNA accumulation profile during the heat time course allowed us to divide miRNAs into four groups - (1) high level after 1h of stress and consecutive decrease of miRNA level when plants were exposed to 3 and 6h of stress; (2) low level of mature miRNA that consecutively increased when plants were exposed to 3 and 6h of stress; (3) increased miRNA accumulation after 1h that does not change during heat stress time course; and (4) miRNA level is increased/decreased only in selected time points of stress conditions (Figure 2). Our results indicate that in barley the strongest response to high temperature stress at the level of mature miRNAs is observed after 6h of heat duration.

**Figure 2.**
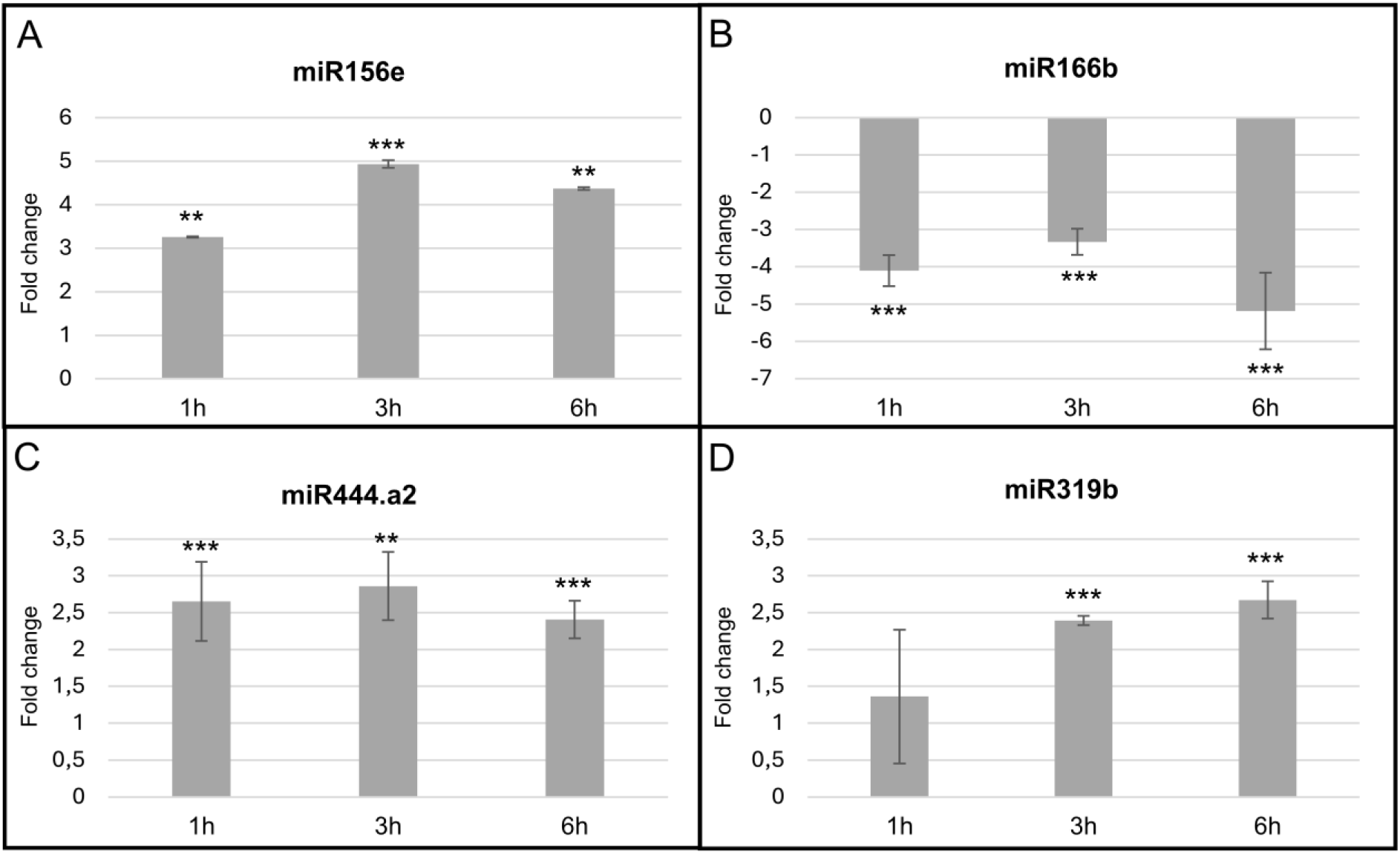
Time course of selected mature miRNAs accumulation. Each graph shows the level of miRNA detected by microtranscriptome analysis in heat-treated barley plants after 1, 3 and 6h of stress treatment. A) miRNAs belonging to group 1 (constant increase of miRNA level); B) miRNAs belonging to group 2 (constant decrease of miRNA level); c) miRNAs belonging to group 3 (miRNA increase after 1h and does not change during heat stress time course); d) miRNAs belonging to group 4 (miRNA level is increased/decreased only in selected time points of stress conditions). EDGE (Empirical analysis of DGE) method was used to find significant fold changes in miRNA accumulation (*p ≤ 0.05, **p ≤ 0.01 and *** p ≤ 0.001).

### Both transcriptional and post transcriptional control affect final accumulation of miRNAs in heat stress

We have used previously published RT-qPCR platform to asses changes in expression level of pri-miRNAs of the most abundantly accumulated miRNAs in the stress conditions tested in this study [53, 54]. For 23 pri-miRNAs of highly accumulated miRNAs the primers were not available at MiREx 2.0 platform. Therefore we have mapped the miRNAs to the barley genome [48] without any substitutions allowed. The genomic sequences surrounding identified miRNA loci were modeled into hairpin structure and primers were designed where at least one of them was anchored within the pre-miRNA structure. The list of pre-miRNA sequences, genomic loci and pre-miRNA secondary structures are listed in Table S2. Altogether, the expression level profiles of 71 pri-miRNAs were assessed. The pri-miRNA expression levels were categorized as upregulated, unchanged and downregulated. As in the case of mature miRNAs one hour of heat stress mildly affected pri-miRNAs expression (six pri-miRNAs were upregulated, five were downregulated, while 60 were unchanged) [Table 2]. Out of these 60 unchanged pri-miRNAs, 38 remained unaffected by the 3 or 6h of heat stress treatment [Figure 3]. Three and six hours of heat stress induced expression of 23 and 30 pri-miRNAs, decreased level of 16 and 11 pri-miRNAs, while the levels of 32 and 30 remained unaffected, respectively [Table 2]. As majority of the up- and down-regulated pri-miRNAs were common for 3h and 6h heat treatment we conclude that already 3h of heat represents severe stress in terms of pri-miRNAs accumulation at 2-week developmental stage [Figure 3].

**Table 2.** Overall changes in pri-miRNA expression during heat stress treatment. 3h and 6h heat stress affects expression of similar number of pri-miRNAs. ↑ - upregulated, = - unchanged, ↓ - downregulated.

| heat 1h |  |  | heat 3h |  |  | heat 6h |  |  |
| --- | --- | --- | --- | --- | --- | --- | --- | --- |
| ↑ | = | ↓ | ↑ | = | ↓ | ↑ | = | ↓ |
| 6 | 60 | 5 | 23 | 32 | 16 | 30 | 30 | 11 |

**Figure 3.**
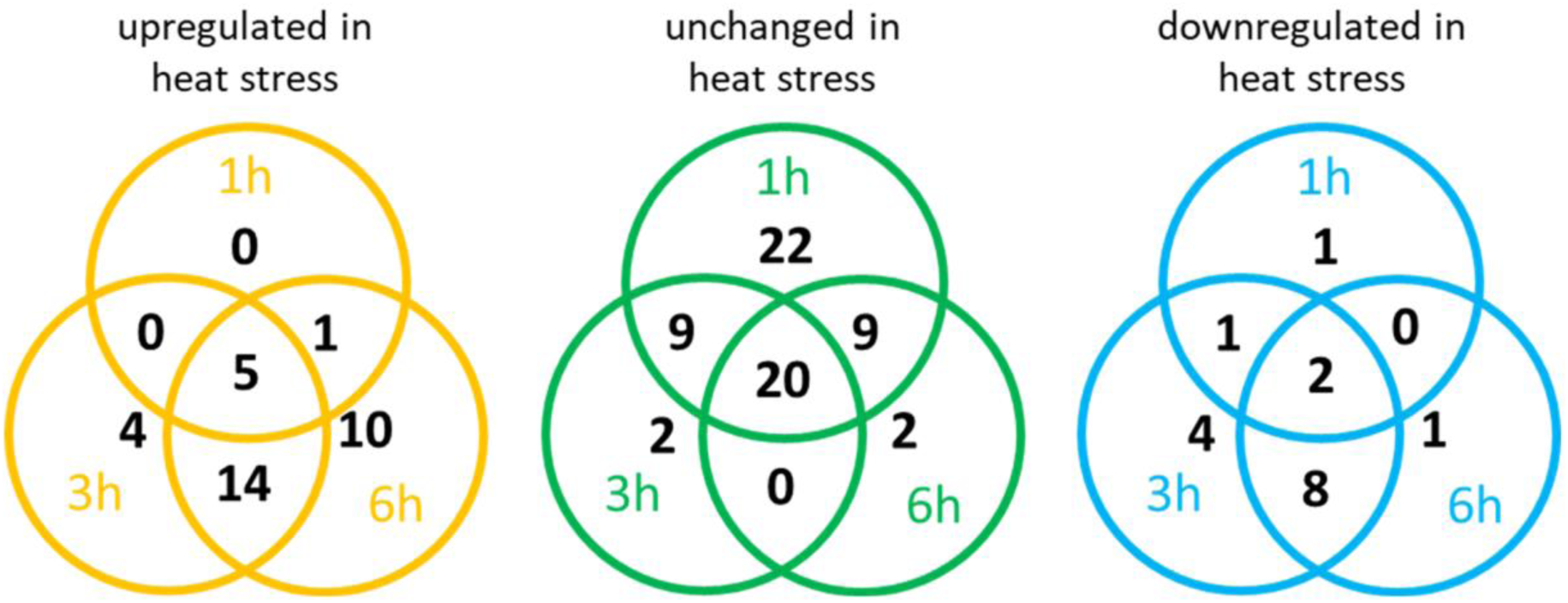
Venn diagrams showing the relations between upregulated, unchanged and downregulated pri-miRNAs during heat stress time course. Most of the pri-miRNAs affected by heat stress are the same for 3h and 6h of stress.

Mature miRNAs expression fluctuations during abiotic stress are not always reflected by the equivalent changes in pri-miRNA levels [9]. The expression profiles altered by heat stress of the 71 tested pri-miRNAs were compared to the accumulation of 62 conserved and three barley specific, cognate miRNAs [Table S4, Table S5]. The correlation between accumulation of miRNAs and the level of their cognate pri-miRNAs can be divided into three groups [Table 3]. The first group are pri-miRNA/miRNA pairs where expression of the pri-miRNA reflects the level of the cognate miRNA. These are 54, 30 and 40 pri-miRNA/miRNA pairs in 1h, 3h and 6h of heat stress, respectively. Out of these pairs, 53 in 1h and 21 in 3 and 6h of heat stress do not change their level. The level of one pri-miRNA/miRNA pair is downregulated in 1h of heat stress, seven and nine are downregulated in 3h and 6h of heat stress, respectively. Two and ten upregulated pri-miRNA/miRNA pairs were detected in 3h and 6h of heat stress. The second group are the pri-miRNA/miRNA pairs where pri-miRNA level is lower when compared to the cognate miRNA level. These are seven, 14 and eight pri-miRNA/miRNA pairs in 1h, 3h and 6h of heat stress, respectively. The third group are the 13, 33 and 34 pri-miRNA/miRNA pairs where the pri-miRNA expression is higher than the cognate miRNA in the respective heat treatments. Therefore, during severe heat stress (3h and 6h of heat) the pri-miRNAs are mostly upregulated in comparison to their miRNA levels. In the case of the second and third group posttranscriptional control of miRNA level during pri-miRNA maturation is postulated while transcriptional control of miRNA level during heat stress takes place in the first group of pri-of miRNA/miRNA pairs. Table S4 presents 28 pri-miRNA/miRNA pairs that likely undergo transcriptional control of miRNA level. Table S5 presents 43 pri-miRNAs where cognate miRNAs accumulate in non-correlated manner suggesting posttranscriptional control of miRNA level. Selected examples of pri-miRNAs and their cognate miRNAs levels during heat stress treatment are presented in Table 4.

**Table 3.** Summary of changes in accumulation of pri-miRNAs in relation to their cognate miRNAs during heat stress treatment.

| pri-miRNA - miRNA relation |  | 1h heat | 3h heat | 6h heat |
| --- | --- | --- | --- | --- |
| pri-miRNA=miRNA | pri-miRNA=, miRNA = | 53 | 21 | 21 |
|  | pri-miRNA↓, miRNA↓ | 1 | 7 | 9 |
|  | pri-miRNA↑, miRNA↑ | 0 | 2 | 10 |
| pri-miRNA<miRNA | pri-miRNA=, miRNA↑ | 3 | 3 | 2 |
|  | pri-miRNA↓, miRNA↑ | 1 | 0 | 0 |
|  | pri-miRNA↓, miRNA= | 3 | 11 | 6 |
| pri-miRNA>miRNA | pri-miRNA↑, miRNA= | 6 | 20 | 23 |
|  | pri-miRNA↑, miRNA↓ | 0 | 5 | 4 |
|  | pri-miRNA=, miRNA↓ | 7 | 8 | 7 |

**Table 4.**
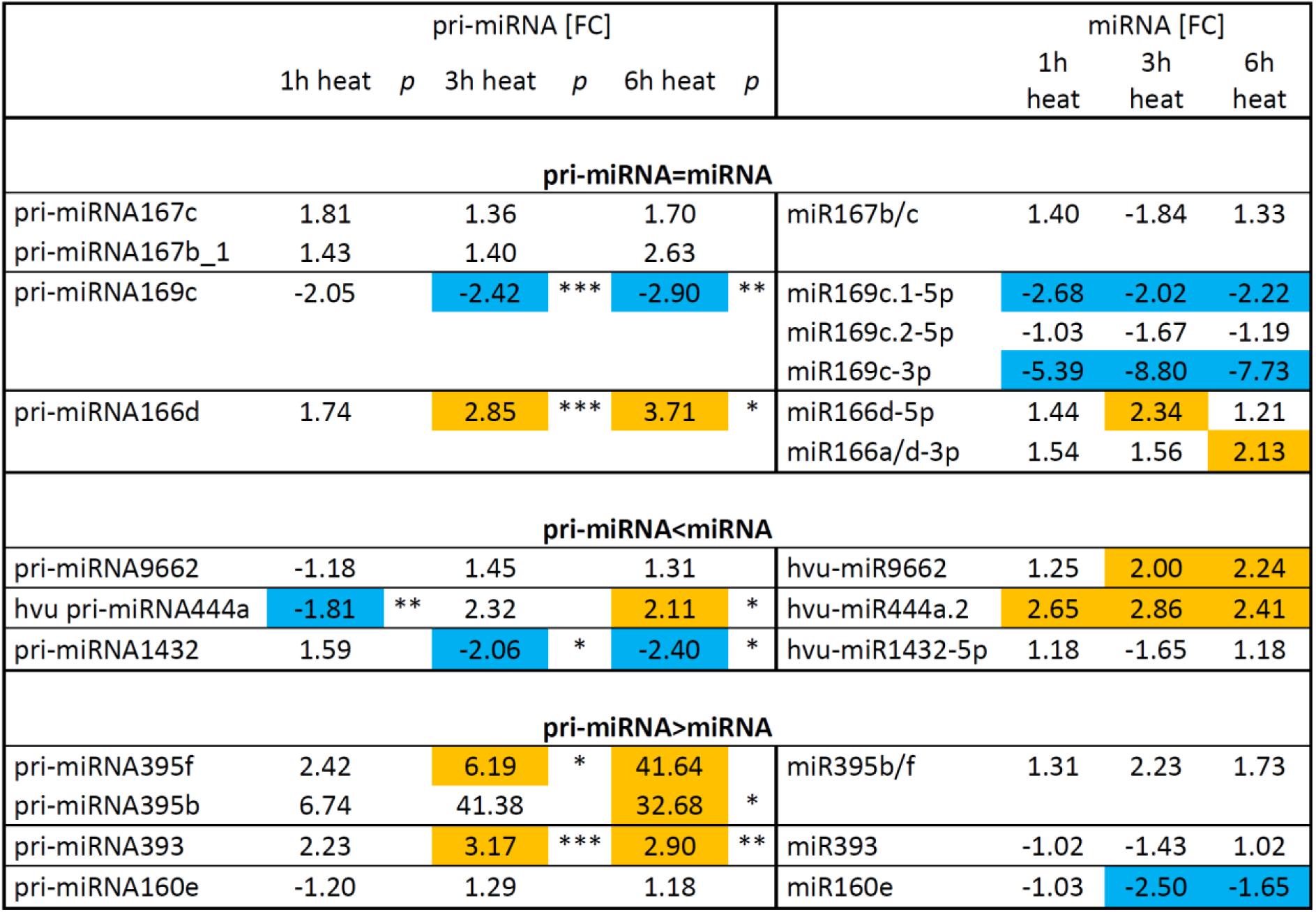
The level of selected barley pri-miRNAs and their cognate miRNAs differently accumulated during heat stress. Date marked in orange show upregulation of pri-miRNA/miRNA while data marked in blue show downregulation of pri-miRNA/miRNA levels. pri-miRNAs and miRNAs are grouped by miRNA families. *P* – pval * ≤0.05; ** ≤0.01; *** ≤0.001. FC, fold change.

### Identification of miRNA-target modules responsive to heat stress

Degradome data obtained for 23-day-old plants were used to identify target mRNAs for conserved miRNAs significantly changed during heat stress [45, 46]. We identified target mRNAs for conserved barley miRNAs (Table S6). For selected target mRNAs their expression level was estimated by RT-qPCR and correlated with the level of mature miRNA (Figure 4). miR319b.1-5p targets transcript of *TCP family transcription factor 4* gene (HORVU2Hr1G060120). miR319b.1-5p is upregulated in heat, while its target is downregulated significantly after 1h of heat, and remained downregulated after 3 and 6h of heat stress. For miR444.a2 significant upregulation was observed after 1h, 3h 6h of heat stress. We observed significant downregulation of its target mRNA encoded by *MADS57* gene (HORVU6Hr1G073040) after 1h of stress. In the case of miR396a we observed its decreased level after 1h, however it is gradually upregulated after 3 and 6h of heat *stress*. miR396a targets mRNA encoded by *Growth-Regulating Factor 4 (GRF4)* gene (HORVU0Hr1G016610) and its level is upregulated upon heat stress, significantly after 1h of stress, however it remains upregulated after 3 and 6h of stress, but not significantly.

**Figure 4.**
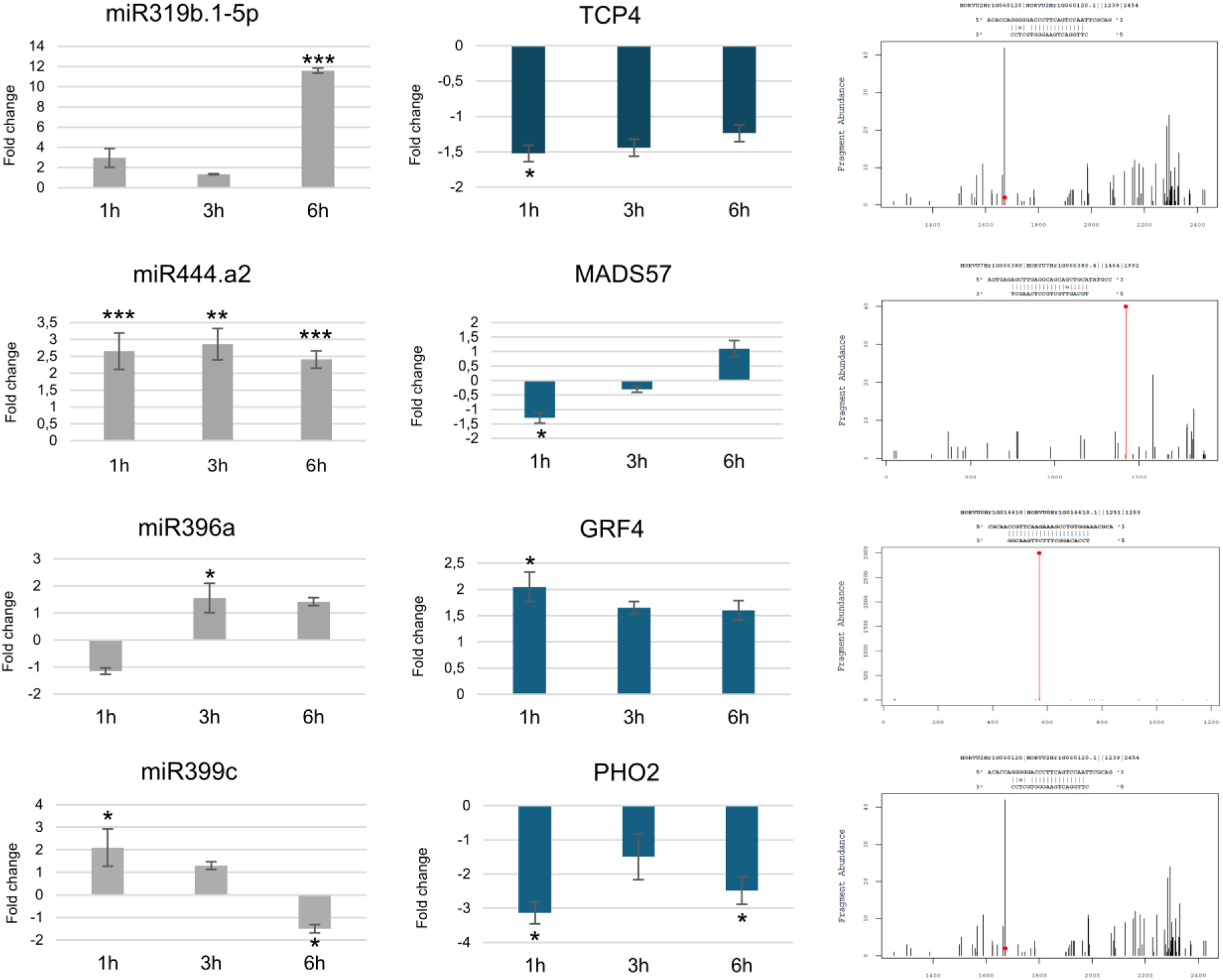
Selected miRNA-mRNA target modules responsive to heat stress treatment (1, 3, and 6h). Left panels represent fold change in miRNA accumulation during heat; middle panels represent target mRNA levels during heat; right panel represent degradome data for miRNA-targeted mRNAs. OY axis shows number of mRNA fragments abundance, while OX axis shows mRNA length in nucleotides. Red line with dot represents miRNA-guided cleavage site in target mRNA. P value - * ≤0.05; ** ≤0.01; *** ≤0.001. TCP4 - TCP family transcription factor 4; MADS57 – MYB transcription factor 57; GRF4 - Growth-Regulating Factor 4; PHO2 - ubiquitin-conjugating E2 enzyme.

The level of miR399c was significantly upregulated after 1h of stress, but in the next hours of stress duration its level was significantly downregulated. The level of mRNA target encoded by *ubiquitin-conjugating E2 enzyme PHO2* gene (HORVU1Hr1G085570) was significantly downregulated during whole heat stress treatment.

### Identification of novel barley miRNA-mRNA target modules

For some miRNAs up to 6 different target mRNAs were identified, usually encoding proteins belonging to the same protein family. Among miRNAs guiding larger number of identified target mRNAs were miR160, 166, and 9662 (Figure S1). miR160 targets mRNAs encoding auxin responsive factors (ARFs). Three barley ARF mRNAs guided for cleavage by miR160a were already published [35] while we identified two novel mRNAs encoding ARF10 (HORVU2Hr1G089670) and ARF16 (HORVU6Hr1G058890) as targets of miR160e (Figure S1A). Six *MIR160* genes were identified in barley (miRbase, release 22.1; [52]. Individual miR160 family members respond to heat stress by increased, decreased or unchanged accumulation. This can be mirrored by their mRNA target levels. The level of miR160a is downregulated after 3h and 6h of heat stress treatment while the expression level of its targets remain unchanged. Only ARF16 mRNA exhibits significant upregulation after 6h of heat stress (Figure S1A). Analysis of IAA auxin indeed shows that auxin level is not affected by heat (Figure S1A).

Barley miR166 family members are known to guide for cleavage mRNAs encoded by *Phavoluta*, *Revoluta* and *Phabulosa* genes as well as mRNA encoding HOX9 transcription factor [35]. We identified two additional transcription factors belonging to HOX TF family as targets for miR166 (HORVU0Hr1G010250 and HORVU3Hr1G026990). MiR166 family represents one of the largest miRNA families in plants with at least seventeen *MIR166* genes encoded in barley (miRbase, release 22.1; [52]. Individual miR166 family members respond to heat stress by increased, decreased or unchanged accumulation. This can be mirrored by their mRNA target levels (Figure S1B). The level of miR166a remains unchanged after heat stress while the expression level of its target mRNAs is slightly decreased, however not significantly (Figure S1B). Two novel target transcripts were also identified for miR9662 encoding proteins belonging to Mitochondrial transcription termination factor family mTERF (HORVU5Hr1G109600 and HORVU7Hr1G040960). Although miR9662 is upregulated in heat stress all identified mTERF targets remain at the same level in heat stress and control plants (Figure S1C).

### miR168/AGO1B target module is affected upon heat stress in barley

Among heat stress responsive barley miRNAs miR168 was detected as upregulated in three time points tested in this study. MiR168 level was increased 1.5 fold after 1h and 3h of heat, and raised up to over 2 fold after 6h of heat stress (Figure 5). Barley miR168 targets mRNA of *Argonaute 1B* gene (*AGO1B*, HORVU7Hr1G007000, [55]) that encodes the component of RISC complex responsible for mRNA slicing. Consequently, *AGO1B* mRNA level was downregulated after 3 and 6h of heat stress compared to control plants (Figure 5). This finding suggests that heat stress may affect miRNA-based posttranscriptional regulation of gene expression in 2-week old barley.

**Figure 5.**
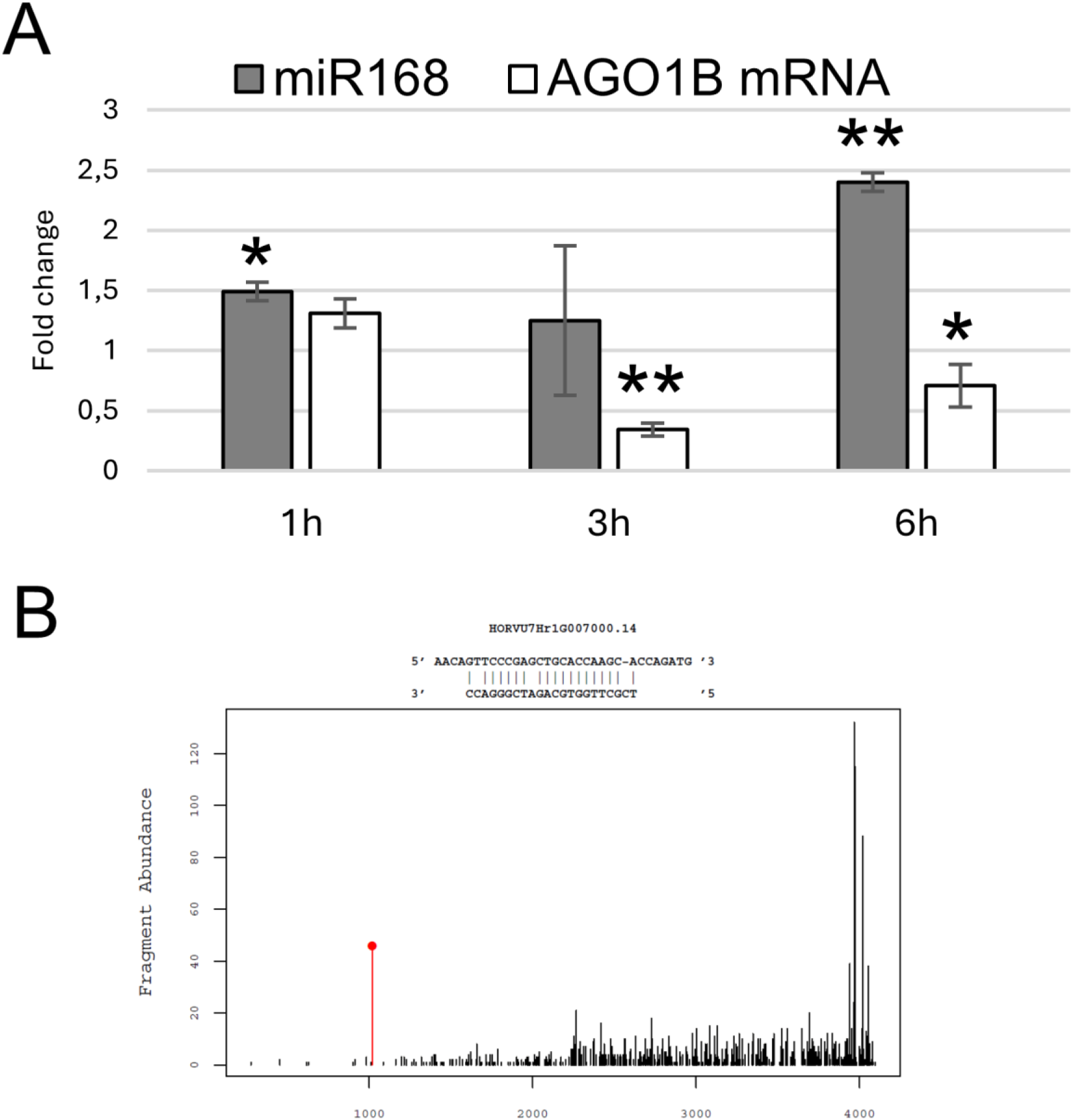
Heat stress affects miR168/AGO1B target module in barley. (A) The level of mature miRNA168 (gray bars) and its target mRNA *AGO1B* (white bars) presented as fold change of accumulation in stress-treated plants compared to control plants. (B) T-plot of *AGO1B* mRNA target, the miRNA driven cleavage site is marked by the red peak. The significance of mRNA expression changes was calculated with Student’s t-test (p value - * ≤ 0.05, and ** ≤ 0.01).

## Discussion

Despite its economic importance, there is no comprehensive report on barley miRNA involvement and regulation in response to heat stress at the early stage of barley development. Previously we reported selected heat-responsive miRNAs [35]. In the present study we performed global analysis of miRNAs in response to a short period of heat stress applied in 2-week-old barley plants entering the tillering period. The most prominent changes in miRNA accumulation were found after 3 and 6h of heat stress duration where the levels of 124 and 155 conserved miRNAs were changed, respectively. This suggests that longer period of heat stress conditions is required for more prominent response at the level of mature miRNAs in young barley plants. Interestingly, the most accumulated miR444c* (∼980 normalized reads) in control barley plants was 3-fold downregulated in 1h and 3h heat treated plants. Interestingly, miR444c was slightly upregulated only after 6h of heat stress. Monocot specific miR444c [56, 57] was reported to target rice and barley MADS box transcription factor 27 responsive to nitrogen supply [58–60]. Deregulation of specific miRNAs* level in response to various stress conditions was previously described [61, 62]. Attempts to find putative target for miR444c* in degradome data were unsuccessful. Moreover, MADS27 mRNA does not show potential cleavage site as possible target for miR444c* in degradome data. There are reports on miRNAs and their miRNAs* coordinated regulation in response to stress conditions. For example it was found that miR172b-5p (considered miRNA*) was downregulated while its partner miR172b-3p was upregulated in response to drought stress. Both miRNA and miRNA* were proven to be functional in flowering acceleration and trehalose increase upon drought stress [63]. Our data suggest that both miR444c and miR444c* may function in response to heat stress conditions by regulating biological pathways improving plant fitness in changing environment. However, the role of miR444c* remains elusive.

We found that upon heat stress the most pronounced differences in accumulation were observed for miRNAs belonging to miR156, 159, 169 and 444 families. While miR156, 159 and 169 families are conserved among monocot and dicot plant species [56, 64], miR444 family is specific only for monocot plants [28, 29, 57]. Members of miR156 and miR169 families were also identified as heat stress responsive in Arabidopsis and wheat [37, 65, 66]. Thus selected miR156 and 169 family members response to heat stress is evolutionary conserved. Our previous studies indicated involvement of other miR444 family members in response to nitrogen supply [28]. Here we show that miR444a2 is strongly upregulated in heat stress. Although further studies are required to clarified the biological role of miR444 family members, it is possible that their main role is to shape monocot plant response to environmental cues.

The expression of 71 pri-miRNAs was profiled and compared to the accumulation of cognate miRNAs during applied heat stress conditions. As previously described for drought stressed barley it is difficult to predict mature miRNA accumulation at the basis of pri-miRNA expression profiles [63]. The discrepancies between mature miRNA and pri-miRNA levels were reported also in Arabidopsis, in plethora of abiotic stresses like heat, drought, salinity, copper, cadmium and sulfur [9]. The uncoupling of pri-miRNA levels and miRNA accumulation is a result of miRNA multi step biogenesis pathway. It is possible that the level of miRNA biogenesis factors changes upon heat stress. For example our data show that *AGO1B* mRNA is significantly downregulated in heat stress. This suggests that AGO1B protein is also downregulated. AGO1B is one of the main executors of miRNA activity. Downregulation of its level may strongly affect the level of many miRNAs in the cell [67]. Heat stress promotes chromatin reorganization in Arabidopsis and affects gene expression [68]. We cannot exclude that in barley genome similar chromatin reorganization events take place affecting *MIR* genes expression.

In the case of heat stressed barley plants, lower number of miRNAs is regulated at transcriptional level, while the majority is regulated post transcriptionally. Similar observation was made in heat stressed Arabidopsis [9]. Interestingly accumulation of some miRNAs is regulated in the same way during heat in barley and Arabidopsis. For instance, the family of miRNA166 is strongly upregulated at the level of pri-miRNA, while the originating miRNAs are unchanged or downregulated. Pri-miRNA319 family is upregulated, and the miRNAs and miRNA*s are differentially expressed. Another interesting example is miR160 family, where pri-miRNAs are upregulated, while miRNAs are downregulated in barley and Arabidopsis or sometimes unchanged in Arabidopsis. In both species, the most prominently upregulated are pri-miRNA395 family members, while originating miRNAs are of unchanged level. These miRNA families may play particularly important role in heat stress response in many plant species pointing to their evolutionary conserved mechanism of action. In barley splicing was proven to be a substantial part of posttranscriptional regulation of pri-miRNA160a and pri-miRNA5175a maturation during heat [35]. Importantly, pri-miRNA160a splice isoforms are highly accumulated in heat stress. However, the most prominently accumulated splice variant is characterized by the lack of hairpin structure. These observations may suggest that the ratio between nonfunctional and functional splice isoforms may regulate the accumulation of miRNA in heat. Another interesting observation is the downregulation of miRNA* in the case of upregulated pri-miRNA159a and pri-miRNA319b after 3h and 6h of heat duration. Such events were reported in heat stressed Arabidopsis seedlings and were characteristic for this abiotic stress [9]. In barley, pri-miRNA172b remains at the same level in control and drought conditions while the levels of miR172b-5p and miR72b-3p derived from this precursor are reversely corelated [63]. Therefore, the discrepancies between different miRNAs accumulation originating from common precursor are a part of posttranscriptional regulation of miRNAs expression. There are many positive and negative regulatory proteins interacting with the miRNA biogenesis complex that affect the ratio of pri-miRNA to miRNA [69]. It must be underlined that the structures of miRNA genes and the processing of individual barley pri-miRNAs is still largely unknown. Therefore, further studies are needed to understand microtranscriptomic changes in barley responses to stresses.

Together with mature heat-affected miRNAs and their cognate precursor levels we also analyzed the expression pattern of target mRNAs in response to heat stress. In plants, miR319 was one of the first characterized and conserved miRNA families, which has been demonstrated to target *TCP* genes. TCP transcription factors participate in the regulation of diverse physiological and biological processes, such as phytohormone biosynthesis and signal transduction, branching, leaf morphogenesis flower development and senescence, pollen development, and regulation of the circadian clock in various plants [70, 71]. The level of miR319b.1-5p was significantly increased in barley and showed an inverse correlation with its target *TCP4* mRNA in heat stress. Similar correlation was found in cotton [72], while in Arabidopsis the level of miR319 family members was downregulated upon heat stress [9]. In Arabidopsis the level of miR396 family members was downregulated upon heat stress, however in rice and barley the level of miR396a was upregulated ([73]; this study). miR396a targets mRNA encoded by *GRF4* gene and its level was upregulated in barley as well in rice.

MiR399c/ PHO2 module was extensively studied in phosphate homeostasis and little is known about its behavior in heat stress. MiR399c targets the *PHO2* gene, that plays a key role in adaptation to phosphate availability [74]. Upregulation of miR399 level combined with downregulation of its target mRNA, *PHO2,* after 1h of heat stress confirmed nicely our previous results [75].

In the present study we also identified novel barley miRNA-guided mRNA targets that respond to heat. Among miRNAs guiding novel target mRNAs miR160e, 166a and 9662 were identified. Both, miR160e and 166a belong to conserved, multi member miRNA families that have been involved in response to different abiotic stresses in Arabidopsis [9, 65], wheat [76, 77] and rice [78]. miR160 targets mRNAs encoding auxin responsive factors (ARFs). In addition to previously identified barley ARF mRNAs as targets for miR160a [35] we found two novel mRNAs encoding ARF10 and ARF16 as targets for miR160e. Heat stress caused fluctuations in the level of specific members of miR160 family and their targets. The analyzed auxin (IAA) level remained unaffected. Gray et al. [79, 80] reported auxin level (IAA) increase in 9-day-old Arabidopsis seedlings when heat stress was applied. Further studies are required to understand the regulation of auxin level in response to heat stress in barley.

MiR166 family is often referred as evolutionarily conserved stress biomarker in land plants [81]. However, in our studies members of this family do not exhibit significant changes in the short period of heat stress applied. Earlier studies on barley revealed that members of this family respond later (after 24h) to heat stress [35]. Thus we can conclude that barley miR166 family members can be used as biomarkers of prolonged heat stress. MiR166 family members are known to guide for cleavage mRNAs encoded by *Phavoluta*, *Revoluta* and *Phabulosa* genes as well as mRNA encoding HOX9 transcription factor [35]. Two additionally identified target mRNAs belonging to HOX32 transcription factor family that are known to regulate a variety of developmental processes and abiotic stress responses in plants have been identified [82, 83]. As expected their level did not changed significantly in stress conditions applied in this study. Plant mitochondrial transcription termination factor (mTERF) family regulates organellar gene expression (OGE) and was functionally characterized in diverse species [80]. Firstly characterized in animal mitochondria, mTERF proteins were involved in mitochondrial transcription, translation, and DNA replication [84]. Transcriptomic analysis suggested that many mTERFs may play important roles in barley development and response to abiotic stresses such cold, salt, and metal ion pollution [80]. In barley, the level of *mTERF15* mRNA was shown to be regulated by miR9662 (miRNA hvu-x11) in response to drought [85]. We identified two novel target mRNAs for miR9662 encoding proteins belonging mTERF family (HORVU5Hr1G109600 and HORVU7Hr1G040960). However, further studies are required to learn more about the positive or negative effects mTERF level on barley response heat stress. In plants, an essential part of PTGS mechanism is exhibited by AGO1 proteins [86]. AGO1 together with miRNA are the main components of the miRISC [67]. Importantly, AGO1 expression is regulated by miR168, creating a feedback mechanism assuring strict control of AGO1 action in PTGS [87]. The proper expression of miR168 in Arabidopsis is important regulator of development [67]. The feedback control mechanism of AGO1 is present also in heat stressed barley plants. In barley two AGO1 genes are present, AGO1A and AGO1B [88]. In this and previous studies we show that barley AGO1B is a subject of miR168 directed cleavage [55]. Heat stress induced the accumulation of miR168 in 1h, 3h and 6h heat stress, while the target mRNA of AGO1B is downregulated in 3h and 6h stress. Therefore we show that in barley, AGO1B-driven PTGS is strictly controlled by feedback mechanism in heat stressed barley plants.

Overall, our studies provide new data on miRNAs and their targets in early response of 2-week-old barley plants to heat. Further studies should concentrate on miRNA/mRNA modules that undergo significant changes under heat stress to understand their role in shaping crop plant response to global climate changes.

## Conclusion

This study provided valuable insights into the complex miRNAs and their targets regulation in response to heat stress at the early stage barley development. Comprehensive sRNA analysis showed that the majority of barley miRNAs responds to heat stress after 3h and 6h of heat stress duration indicating that for more prominent response at the level of mature miRNAs longer period of heat stress is required. Besides, when changes in mature miRNA accumulation were compared to their cognate precursor levels it has been shown that majority of heat-affected miRNAs was regulated on post transcriptional level. The identified miRNA-mRNA target modules also demonstrated that wide range of biological processes and pathways, including miRNA biogenesis itself, may be impaired by heat stress in young barley plants.

## Acknowledgments

This work was supported by Narodowe Centrum Nauki (National Science Centre, Poland), grant UMO-2016/21/B/NZ9/00550. AP was supported by grant UMO-2016/23/B/NZ9/00857.

## Data availability statement

The small RNA-seq data that support the findings of this study are available in NCBI’s Gene Expression Omnibus, GEO and are accessible through GEO Series accession number GSE272607; https://www.ncbi.nlm.nih.gov/geo/query/acc.cgi?acc= GSE272607.

## Supplementary Tables

Table S1. List of primer sequences used in RT-qPCR reactions.

Table S2. The list of barley pre-miRNA sequences and their respective genomic loci with coordinates. The pre-miRNA stem-loop structures were constructed using FolderVersion1.11 software. MiRNA position in stem-loop structures is marked in color. ΔG – free energy kcal/mol.

Table S3. Barley miRNAs differentially accumulated in heat stress. Small RNA sequences were annotated to miRbase Sequence database (release 22) without mismatches and with strand-specific alignment. For further analysis miRNAs with length of 20-22 nt. with read score more than 10 and pval ≤ 0.05 were selected.

Table S4. Barley pri-miRNAs and cognate miRNAs with correlated changes in their levels under high temperature stress. Data marked in orange show upregulation of pri-miRNA/miRNA while data marked in blue show downregulation of pri-miRNA/miRNA levels. pri-miRNAs and miRNAs are grouped by miRNA families. *P* – pval * ≤0.05; ** ≤0.01; *** ≤0.001. FC, fold change.

Table S5. Barley pri-miRNAs and cognate miRNAs that show no correlation in their level under high temperature stress. Data marked in orange show upregulation of pri-miRNA/miRNA while data marked in blue show downregulation of pri-miRNA/miRNA levels. pri-miRNAs and miRNAs are grouped by miRNA families. *P* – pval * ≤0.05; **≤ 0.01; *** ≤0.001. FC, fold change.

Table S6. List of barley target mRNAs identified for miRNAs most prominently changed during heat stress treatment using degradome data.

## Supplementary Figure

Figure S1. Novel members of conserved target families identified for miRNAs guiding for cleavage larger number of target mRNAs.

Left panels represent T-plots of mRNA targets, the miRNA-driven cleavage site is marked by the red peak; middle panels represent expression of target mRNAs for (A) miR160e, (B) miR166a and (C) miR9662, respectively; right panel in (a) shows the level of IAA auxin (indole-3-acetic acid) in control (white bars) and heat treated plants (grey bars). p value * ≤ 0.05.

