## Supplementary figures and images for "Barley miRNAs and their targets regulation in response to heat stress at the early stage of development"

### Figure S1

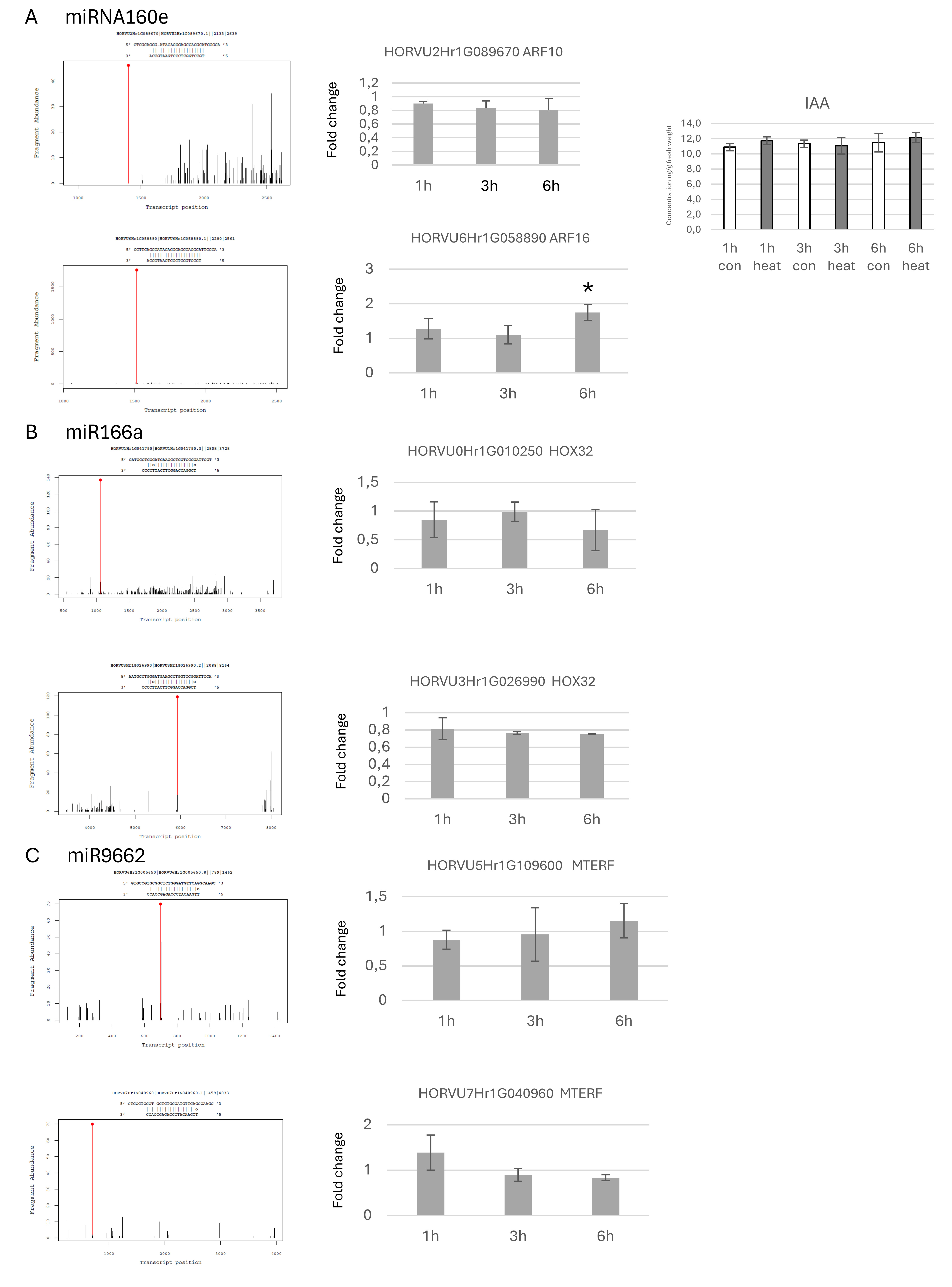
