## Supplementary material for "Barley miRNAs and their targets regulation in response to heat stress at the early stage of development": Table S1

Table S1. List of primer sequences used in RT-qPCR reactions.

| Primer name | Primer sequence (5' --> 3') | Target | usage |
|---|---|---|---|
| OS927 | GGATTGAAGGGAGCTCTGCA | pri-miRNA159a | Fwd RT-qPCR |
| OS928 | AGAGCTTGCACCCAGATCTG | pri-miRNA159a | Rev RT-qPCR |
| HvReTi4F | TGTTTTGGATTTGGTTTGGAG | pri-miRNA159a | Fwd RT-qPCR |
| HvReTi4R | ACCACAAGCCTATCTCCTCGT | pri-miRNA159a | Rev RT-qPCR |
| KK764 | ACCGCTATTTGGATTGAAGG | pri-miRNA159b | Fwd RT-qPCR |
| KK765 | TCTGCAGTAGTTGGTTCCAGAT | pri-miRNA159b | Rev RT-qPCR |
| OS205 | CCAAGCATGACCGTCTCTCT | pri-miRNA160a | Fwd RT-qPCR |
| OS206 | GATCGGGTTACCCTCTACCA | pri-miRNA160a | Rev RT-qPCR |
| KK760 | GCTCCCTGTATGCCACTCAT | pri-miRNA160b | Fwd RT-qPCR |
| KK761 | AGGGAGAGGGTTTGATTTCG | pri-miRNA160b | Rev RT-qPCR |
| KK758 | TTGACGATCGACCTAGTGAAGA | pri-miRNA160c_1 | Fwd RT-qPCR |
| KK759 | AGGAAGAATGCAATAGAAGACGA | pri-miRNA160c_1 | Rev RT-qPCR |
| KK756 | GCTTGATCAGGTAGTCGTGGT | pri-miRNA160c_2 | Fwd RT-qPCR |
| KK757 | TCAAGATGGGATCGAAGATAGA | pri-miRNA160c_2 | Rev RT-qPCR |
| KK762 | CTGAATGCCATCCGAGAAG | pri-miRNA160e | Fwd RT-qPCR |
| KK763 | GAAAGTATAGTAAAGGAAGGAGGAGGA | pri-miRNA160e | Rev RT-qPCR |
| OS923 | TGGATGGATGGATGGTGCAA | pri-miRNA160f | Fwd RT-qPCR |
| OS924 | TTCTCGGATGGCATTCAGGG | pri-miRNA160f | Rev RT-qPCR |
| OS949 | TGCATGAGAGAGGGTGAGGA | pri-miRNA166a | Fwd RT-qPCR |
| OS950 | GCGAGACCTTGAACCAGACA | pri-miRNA166a | Rev RT-qPCR |
| OS913 | CGGCGACCTACTTAGCTAGC | pri-miRNA166c | Fwd RT-qPCR |
| OS914 | GCCAACGTTCCCCACAAAC | pri-miRNA166c | Rev RT-qPCR |
| OS220 | CTGGTTCAAGGTCTCGCTCT | pri-miRNA166d | Fwd RT-qPCR |
| OS221 | CCCCCTTACCCAAGAAACAT | pri-miRNA166d | Rev RT-qPCR |
| HvReTi_47F | GTTGTCTGGTTCAAGGTCTCG | pri-miRNA166e | Fwd RT-qPCR |
| HvReTi_47R | ACATCAACAATCTTTGGCTGG | pri-miRNA166e | Rev RT-qPCR |
| OS216 | GCATGAGAGAGGGTGAGGAA | pri-miRNA166f | Fwd RT-qPCR |
| OS217 | CAGGCGAGAGAAATGGAAGT | pri-miRNA166f | Rev RT-qPCR |
| HvReTi_5F | CAGCGTCATCTTCTTCGTTTC | pri-miRNA166n | Fwd RT-qPCR |
| HvReTi_5R | AGGTGGAGCTCACAAGAACAC | pri-miRNA166n | Rev RT-qPCR |
| HvReTi_52F | GCTGCCAGCATGATCTAACTC | pri-miRNA167b | Fwd RT-qPCR |
| HvReTi_52R | CCCAACAGGGAAAGAGTGAA | pri-miRNA167b | Rev RT-qPCR |
| HvReTi_41F | TTGATGGGTAGATCAAGGTGC | pri-miRNA167b | Fwd RT-qPCR |
| OS203 | TGGGATTTTAGGGTTTTCGTT | pri-miRNA167b | Rev RT-qPCR |
| KK738 | TTCTTGGTTAGGATGGAAATGC | pri-miRNA167c | Fwd RT-qPCR |
| KK739 | AGGCCATACTGCAAACACATAG | pri-miRNA167c | Rev RT-qPCR |
| KK823 | AGCTATGCTGCATGTCGATCT | pri-miRNA167d | Fwd RT-qPCR |
| KK824 | GTTTCCTCCCGTAGATCTTGC | pri-miRNA167d | Rev RT-qPCR |
| KK939 | CCAAATTTCAACACTAACATCAGG | pri-miRNA169a | Fwd RT-qPCR |
| KK940 | CCTAGCATGAGGAGCAGGAG | pri-miRNA169a | Rev RT-qPCR |
| OS909 | CAAAGCAGTCAGGGTAGGGG | pri-miRNA169c | Fwd RT-qPCR |
| OS910 | ATCCCGTTCTCTGCCCCTTA | pri-miRNA169c | Rev RT-qPCR |
| OS915 | GGTGAAGCCTTGCATGGAGA | pri-miRNA169d | Fwd RT-qPCR |
| OS916 | TGGTGAGATGGATCAGTTAGCA | pri-miRNA169d | Rev RT-qPCR |
| KK941 | GCTGCAAGGGCCTTATCTCT | pri-miRNA169e | Fwd RT-qPCR |
| KK942 | TGAAGAGACAGCCTAGCATGG | pri-miRNA169e | Rev RT-qPCR |
| KK945 | TGCAGTAGCAGAGAGCAAGC | pri-miRNA169g | Fwd RT-qPCR |
| KK946 | TCAGTATGCATTGATTTAATAAAGAGC | pri-miRNA169g | Rev RT-qPCR |
| OS959 | TTCCATGATAGGCGGTCACC | pri-miRNA169h | Fwd RT-qPCR |
| OS960 | ACCTGTCAGCTGCAATTCAA | pri-miRNA169h | Rev RT-qPCR |
| OS919 | AGGGAGCAGGAGGAAGAAGA | pri-miRNA169i | Fwd RT-qPCR |
| OS920 | CATCCACAGGCAAGTCATCCT | pri-miRNA169i | Rev RT-qPCR |

| | | | |
|---|---|---|---|
| KK943 | CTGCAAGGGCCTTATCTCTG | pri-miRNA169i | Fwd RT-qPCR |
| KK944 | ACACCCCAAGGCCAATTAAGCC | pri-miRNA169i | Rev RT-qPCR |
| OS222 | GTGAGGTTCAATCCGATGCT | pri-miRNA171a | Fwd RT-qPCR |
| KK667 | CTGCCATGGAGATGTAGTTGG | pri-miRNA171a | Rev RT-qPCR |
| OS224 | CAACCAGTCAAGGCAAGGTT | pri-miRNA171b | Fwd RT-qPCR |
| OS225 | GATGGCTTGCGACTCAATTT | pri-miRNA171b | Rev RT-qPCR |
| KK672 | TCAGAGCTCTTGTCATTTCATCA | pri-miRNA171c | Fwd RT-qPCR |
| KK673 | AGTTGCGACGCTAGCTTAATTT | pri-miRNA171c | Rev RT-qPCR |
| KK670 | AGAAAACAAAGACGAAATGAAATTG | pri-miRNA171d | Fwd RT-qPCR |
| KK819 | ATGCAGGAGATCGATGTTGAC | pri-miRNA171d | Rev RT-qPCR |
| HvReTi_12F | CTCTCCTCCTTGCGGGTTGAT | pri-miRNA171e | Fwd RT-qPCR |
| HvReTi_18R | GCGGAGGAGCTAAGCTAGGTA | pri-miRNA171e | Rev RT-qPCR |
| KK674 | AAGAAGAAGACGACATGCTGGTA | pri-miRNA171h | Fwd RT-qPCR |
| KK675 | AGGGCTTCTCTGAAGGAAGAATA | pri-miRNA171h | Rev RT-qPCR |
| HvReTi_26F | CGCCATGGAAATATATGAGAGG | pri-miRNA172b | Fwd RT-qPCR |
| HvReTi_26R | GATGTGAATCTTGGTGGTGCT | pri-miRNA172b | Rev RT-qPCR |
| HvReTi28_F | AATTAGCTGCCGACTCATTCA | pri-miRNA319b | Fwd RT-qPCR |
| HvReTi28_R | CGAAGAAACGAGCAGATCAAG | pri-miRNA319b | Rev RT-qPCR |
| OS951 | CCATTCGGGTGATTAGCTAGCT | pri-miRNA393 | Fwd RT-qPCR |
| OS952 | TGACGGAGGGAGATCGATCA | pri-miRNA393 | Rev RT-qPCR |
| KK588 | GCCATCAGAGAGCGAGAGAG | pri-miRNA394_1 | Fwd RT-qPCR |
| KK589 | GATTCAGGAGGAGGTGGACA | pri-miRNA394_1 | Rev RT-qPCR |
| OS963 | GGAGGGGAGAGAGCTTGAAG | pri-miRNA394_2 | Fwd RT-qPCR |
| OS964 | TCAGGAGGAGGTGGACAGAA | pri-miRNA394_2 | Rev RT-qPCR |
| KK586 | CCAAGCATTTAGCCGACATT | pri-miRNA395b | Fwd RT-qPCR |
| KK587 | TGCTTGGTTACACCGAGAGTT | pri-miRNA395b | Rev RT-qPCR |
| KK582 | CGGAGTTCCTTTGATGCACT | pri-miRNA395f | Fwd RT-qPCR |
| KK583 | GCTAGAAGTAACCTCCCGGTAA | pri-miRNA395f | Rev RT-qPCR |
| KK572 | TGCTCTCCACAGGCTTTCTT | pri-miRNA396a | Fwd RT-qPCR |
| KK573 | GAACCATCAACAAGGCAAGG | pri-miRNA396a | Rev RT-qPCR |
| KK708 | GATCTCTCGTTCAAGCTAGTCCA | pri-miRNA396c | Fwd RT-qPCR |
| OS182 | GAGGAGGGGAGAGGAAACAG | pri-miRNA396c | Rev RT-qPCR |
| KK578 | GGTGCTGGATGCTGATTTCT | pri-miRNA396e | Fwd RT-qPCR |
| KK579 | TTGGCCTCTCTCTGCAATTT | pri-miRNA396e | Rev RT-qPCR |
| KK911 | AGTGGACGTTGGAGGAGGAG | pri-miRNA396h | Fwd RT-qPCR |
| KK912 | CTCCAAGTACTGCGAGAAGCA | pri-miRNA396h | Rev RT-qPCR |
| OS907 | CCATGATTCCTGCTTGCGTC | pri-miRNA397a | Fwd RT-qPCR |
| OS908 | TCATCAACGCTGCACTCAAC | Fpri-miRNA397a | Rev RT-qPCR |
| KK568 | TCGCTGCAGGTCAGTTACAA | pri-miRNA398f | Fwd RT-qPCR |
| KK569 | TGTGGAATCGAGGAGAGGAG | pri-miRNA398f | Rev RT-qPCR |
| KK907 | CAGAGGGATGGATGTGGATT | pri-miRNA399d_1 | Fwd RT-qPCR |
| KK1012 | GCAAATCTCCTTTGGCAGAC | pri-miRNA399d_1 | Rev RT-qPCR |
| OS230 | TGGCGGTGATTAGAAAGTGA | pri-miRNA399d_2 | Fwd RT-qPCR |
| OS231 | GCAAATCTCCTTTGGCAGAC | pri-miRNA399d_2 | Rev RT-qPCR |
| OS228 | TGTTGGCCCCTCTTACTTTG | pri-miRNA399g_1 | Fwd RT-qPCR |
| KK721 | AGAGAGACCAACAATGGATGC | pri-miRNA399g_1 | Rev RT-qPCR |
| OS226 | GGTGGCGAGTGAGATAGGTG | pri-miRNA399g_2 | Fwd RT-qPCR |
| OS227 | AAGACCAACAAGAGCGCAAC | pri-miRNA399g_2 | Rev RT-qPCR |
| KK558 | GGGTGAGAATCACAGTGCAG | pri-miRNA399h | Fwd RT-qPCR |
| KK559 | TGCAAATAGCAGTGTGCCATA | pri-miRNA399h | Rev RT-qPCR |
| OS901 | GCATCAGGTGAGGTGGTGAA | pri-miRNA408 | Fwd RT-qPCR |
| OS902 | CATCCCTTGCTCTGCTCCAT | pri-miRNA408 | Rev RT-qPCR |
| KK357 | ACCTGCAAGCAAGACGCAAAAT | pri-miRNA444a | Fwd RT-qPCR |
| KK406 | GTTTTTTTCTGATAGCTTTGGCAAAG | pri-miRNA444a | Rev RT-qPCR |
| KK405 | ACTTTCAAGGAAGTTACAAAATC | pri-miRNA444b | Fwd RT-qPCR |

| | | | |
|---|---|---|---|
| KK367 | TTCCTATTTCACATTGCGGTC | pri-miRNA444b | Rev RT-qPCR |
| APO469 | AAGTGGAGGCGGCAAGCTA | pri-miRNA444c | Fwd RT-qPCR |
| APO470 | CGGTATGGCCCGAAAACGAT | pri-miRNA444c | Rev RT-qPCR |
| OS903 | CCGTGTCTCCATGCGATGAT | pri-miRNA528 | Fwd RT-qPCR |
| OS904 | GAGTGTGGGCGACGAACTTA | pri-miRNA528 | Rev RT-qPCR |
| OS905 | TCTCCACGCGATGATGATGG | pri-miRNA528 | Fwd RT-qPCR |
| OS906 | TTTCAAGAGAGTGGGCGACG | pri-miRNA528 | Rev RT-qPCR |
| OS953 | TGGCTAGGCTCTCATTCCCT | pri-miRNA827 | Fwd RT-qPCR |
| OS954 | CGGATGGCAGATGGAACGAT | pri-miRNA827 | Rev RT-qPCR |
| KK682 | CTGCCGGTGCCTATAACATT | pri-miRNA1130 | Fwd RT-qPCR |
| KK683 | AAGATGGAATGAACCATCTGC | pri-miRNA1130 | Rev RT-qPCR |
| HvReTi_11F | TCTTGGTGGTCTTGTGGGTTA | pri-miRNA1432 | Fwd RT-qPCR |
| HvReTi_11R | AAACGGATACATCATGGCCTA | pri-miRNA1432 | Rev RT-qPCR |
| KK855 | TGACAAACCAAAATCCGTCAT | pri-miRNA5048b | Fwd RT-qPCR |
| KK856 | ATGGATATGCATGTGGGACAA | pri-miRNA5048b | Rev RT-qPCR |
| KK690 | AAAACTGATCACTATTACTCCCTGTG | pri-miRNA5049b | Fwd RT-qPCR |
| KK691 | TTCCTCAGAGTGTTCGTGATGT | pri-miRNA5049b | Rev RT-qPCR |
| KK726 | CCCCCATGTCTGTCTGTCTT | pri-miRNA5049e | Fwd RT-qPCR |
| KK727 | GCGCCCTTCCCTAAATTCT | pri-miRNA5049e | Rev RT-qPCR |
| OS969 | TGCTGCTGGGTGGTCTTTAG | pri-miRNA5051 | Fwd RT-qPCR |
| OS970 | AGATGACTCGCCCAAAACCC | pri-miRNA5051 | Rev RT-qPCR |
| OS973 | TTGCCCTGATCTGACCCTTG | pri-miRNA5052 | Fwd RT-qPCR |
| OS974 | CTTGTGGCGGTAGGATGTGT | pri-miRNA5052 | Rev RT-qPCR |
| OS917 | CCATGGGGTAGTAGCTTCACC | pri-miRNA5168 | Fwd RT-qPCR |
| OS918 | CTTGAACCAGACAACAACCCC | pri-miRNA5168 | Rev RT-qPCR |
| OS955 | AAAACACGTGGAAGAATATGCAA | pri-miRNA5200 | Fwd RT-qPCR |
| OS956 | CACATCTTGTCTTTTGTCGGCG | pri-miRNA5200 | Rev RT-qPCR |
| OS961 | TTCACGCAGAAGGAGATCGA | pri-miRNA9662 | Fwd RT-qPCR |
| OS962 | TGAGGATAAACACGGTATTGGGG | pri-miRNA9662 | Rev RT-qPCR |
| OS981 | TATGCTTTCACCGTCGATGC | pri-miRNAx9b | Fwd RT-qPCR |
| OS982 | GCAAACGAGAAATTGGCCCA | pri-miRNAx9b | Rev RT-qPCR |
| OS987 | CAAGCAATTCAAGCATCCAA | pri-miRNAx13 | Fwd RT-qPCR |
| OS988 | GGATCAGAGAGAGGCTGCAC | pri-miRNAx13 | Rev RT-qPCR |
| APO387 | CGTGACGCTGTGTTGCTTGT | ADP | Fwd RT-qPCR |
| APO388 | CCGCATTCATCGCATTAGG | ADP | Rev RT-qPCR |
| Target TCP | ACCGAGGACCAGGACCAG | Target TCP | Fwd RT-qPCR |
| Target TCP | CCGTGTAGACCTTGCTGTGG | Target TCP | Rev RT-qPCR |
| Target MADS57 | TGCAAGAAAGCCACAAGCAA | MADS57 | Fwd RT-qPCR |
| Target MADS57 | TGCGTAAACTCATTTCGAGC | MADS57 | Rev RT-qPCR |
| Target GRF4 | TATTGATGTGCCGCTCACAATAC | GRF4 | Fwd RT-qPCR |
| Target GRF4 | CCTCTTTCGCATTCTTCCTTG | GRF4 | Rev RT-qPCR |
| Target PHO2 | GTTCTTCTGGAAGCTCCCTTG | PHO2 | Fwd RT-qPCR |
| Target PHO2 | GATTTGCCCCGAACTATCG | PHO2 | Rev RT-qPCR |
| Target AGO1B | GCGCTCATCATGGTTAGGAA | AGO1B | Fwd RT-qPCR |
| Target AGO1B | CACGGCCACCTTGTTGA | AGO1B | Rev RT-qPCR |
| Target ARF10 | CCGTCTACTACTTCCCGCAG | ARF10 | Fwd RT-qPCR |
| Target ARF10 | GGACGAGGCGGATCTTGG | ARF10 | Rev RT-qPCR |
| Target ARF16 | AGCCGTTCGTTGTGAGAGTT | ARF16 | Fwd RT-qPCR |
| Target ARF16 | TCGGGAGAGCACCAACGG | ARF16 | Rev RT-qPCR |
| Target HOX32 | TCGTGTGGCTTTTTCTTGTG | HOX32 | Fwd RT-qPCR |
| Target HOX32 | GTAGACGAGACGCGACACC | HOX32 | Rev RT-qPCR |
| Target HOX32 | ACCCCAAGCAGATCAAGGTC | HOX32 | Fwd RT-qPCR |
| Target HOX32 | ATGAGCAGCTTGTTCATGG | HOX32 | Rev RT-qPCR |
| Target mTERF | TCGGCGAGGAGTATATCACA | mTERF | Fwd RT-qPCR |
| Target mTERF | ACAATGTCAGCACCATTGGA | mTERF | Rev RT-qPCR |

| | | | |
|---|---|---|---|
| Target mTERF | GCGTCGTCTCTCATCTCCTC | mTERF | Fwd RT-qPCR |
| Target mTERF | GTAGTCCTCCACGGCGAAG | mTERF | Rev RT-qPCR |
