## Supplementary material for "Barley miRNAs and their targets regulation in response to heat stress at the early stage of development": Table S2

Table S2. The list of barley pre-miRNA sequences and their respective genomic loci with coordinates. The pre-miRNA stem-loop structures were constructed using FolderVersion1.11 software. MiRNA position in stem-loop structures is marked in color. ΔG – folding free energy kcal/mol.

pre-miRNA159a
>3H dna:primary_assembly
primary_assembly:MorexV3_pseudomolecules_assembly:3H:11949371:11949563:-1
GTTTGGAGGTGGAGCTCCTATCATTCCAATGAAGGGTCTACCGGAAGGGTTTGTGCAGCT
GCTTGTTCATGGTTCCCACTATCCTATCTCCATTAGAACACGAGGAGATAGGCTTGTGGT
TTGCATGATCGAGGAGCCGCTTCGATCCCTCGCTGACCGCTGTTTGGATTGAAGGGAGCT
CTGCATCTTGATC

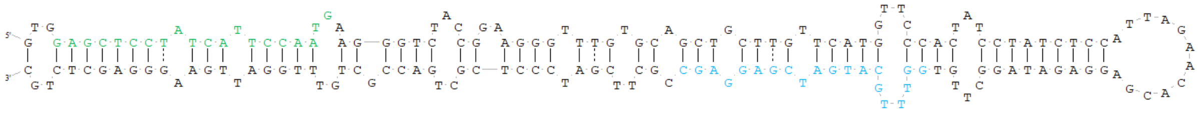

ΔG=-84.6 kcal/mol
pre-miRNA160f
>7H dna:primary_assembly
primary_assembly:MorexV3_pseudomolecules_assembly:7H:158974792:158974893:1
TGGCTGGCGTTTGCCTGGCTCCCTGAATGCCATCCGAGAAGCGCGCCACTGTGGGAGGCG
TCCCTCCTGGTTGGCATTGAGGGAGTCATGCAGGCTCCTCCT

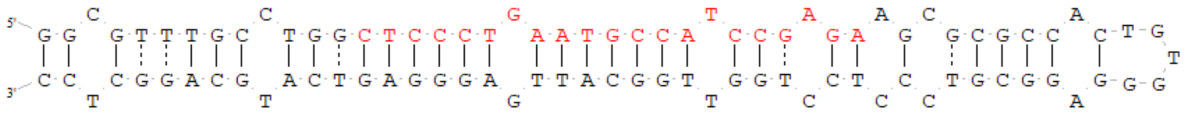

ΔG=-54.1 kcal/mol
pre-miRNA166a
>1H dna:primary_assembly
primary_assembly:MorexV3_pseudomolecules_assembly:1H:335384356:335384498:1
ATATGAAGCTATTTTGCTTCTGGGTGGAATGTTGTCTGGTTCAAGGTCTCGCTCTGAGGT
CTGAGGATGGAAAACTACTTCCATTTCTCTCGCCTGCGAGCGATCTCGGACCAGGCTTCA
TTCCCCTCAGAGATAGCTTCAAC

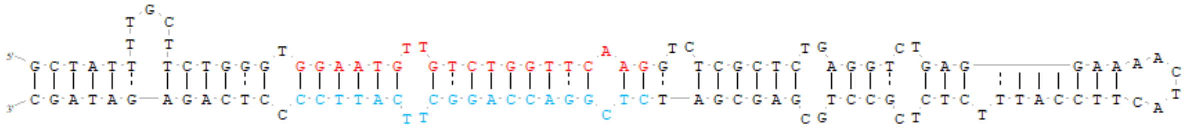

ΔG=-64.1 kcal/mol
pre-miRNA169c
>2H dna:primary_assembly
primary_assembly:MorexV3_pseudomolecules_assembly:2H:604745746:604745908:1
CTTCTGCTGGGTTGCCATACGCTAAGGGGCAGAGAACGGGATGCAGCCAAGGATGACTTG
CCGGCTTCTGGTGTTGGGAGTTCGTAGAGCCTTAAGAATTAGCCGGCAAGCTGTCCTTGG
CTACACCTAGTTCTCTTCTTCTGGTGTTTGGCCCTCAAGCATT

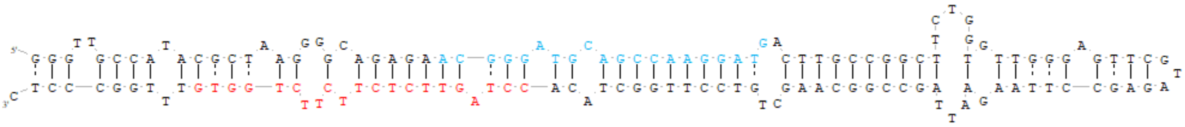

ΔG=-79.5 kcal/mol
pre-miRNA169d
>4H dna:primary_assembly
primary_assembly:MorexV3_pseudomolecules_assembly:4H:60116822:60117052:1
AAGAAGAAGAAGAAGAGGTGAAGCCTTGCATGGAGATGGAGAGCCTCCCTTCATCTGGTA
GCCAAGAATGACTTGCCTATGCAATGCCTCTGCTAACTGATCCATCTCACCATCAATTTG
GGCAAAGTTCTTGTGATGGATGCAGTTTGTGTGGTTGCATGGGTGGGTCTTCTTGGCTAA

CCAGAGTGGCTCTCATCCACCATGCCAGGCCATCCCTCTTCAAGAAACAAA

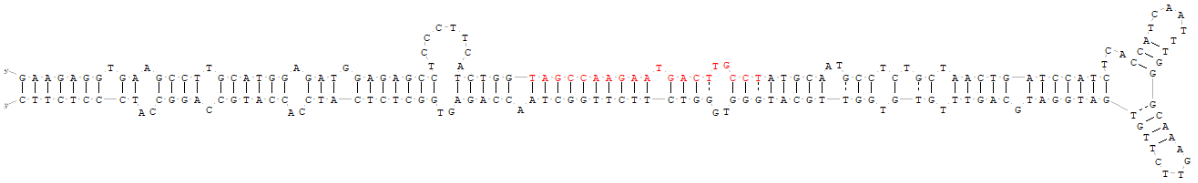

ΔG=-113.8 kcal/mol
pre-miRNA169i
>5H dna:primary_assembly
primary_assembly:MorexV3_pseudomolecules_assembly:5H:489910660:489910885:1
GAAGGGAGTAGTGAGATAGACAAAGAGAGATAGGCCTTGCATGGCGACGAGGGCTTCCTC
TGGTAGCCAAGGATGACTTGCCTGTGGATGTATAGCTTGATATCGATTGCGCTTGCATCC
CTGCTTGTATGTGATCTGTCACTATGTACAACAGGCAGTCTCCTTGGCTAGCCCGAGTGG
CCCTCGTCCTTCATGCCAGCCTCTCATCTTTTCTCAAACCGTAACC

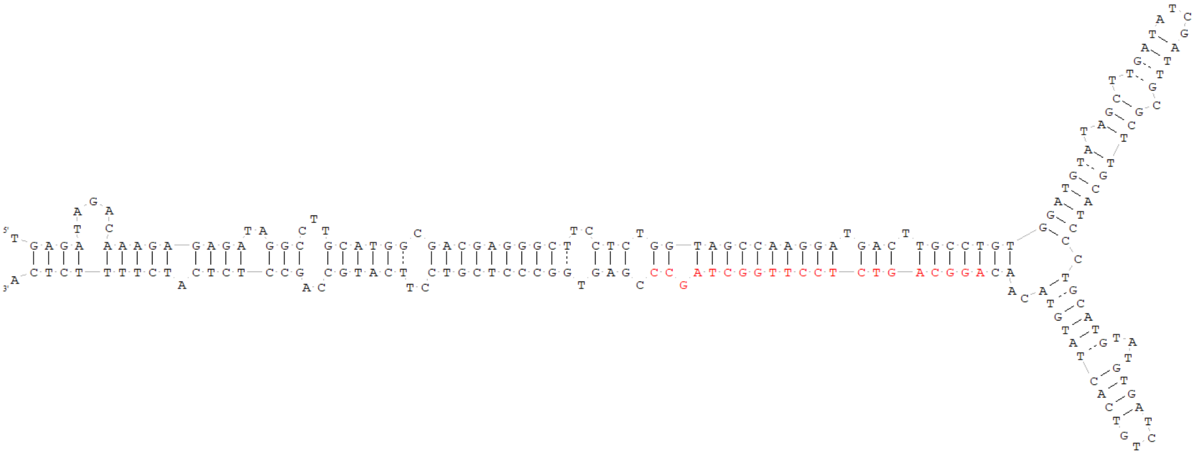

ΔG=-101.1 kcal/mol
pre-miRNA169h
>2H dna:primary_assembly
primary_assembly:MorexV3_pseudomolecules_assembly:2H:440198893:440199122:1
CTGTCACCTCCGGTAGGAAGAAGCTAGGGAGAAGAGGGGCTTTGCATGAGGGCCAGAGCC
TGCTTCATCTGGTAGCCAAGGATGACTTGCCTATATGCTCTTTCAGAAGCCCCGGTTATT
ACATGGATGGTTTCAGATGAGTTCCATGATAGGCGGTCACCTTGGCTAGCCTGAGCAGCT
CTTGTCTGTCATGGAAGGCCTCTCCTTCTCATTCTCTGCCATATCTATCT

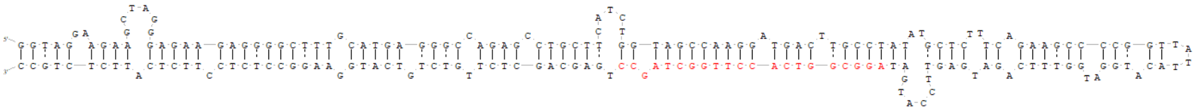

ΔG=-101.9 kcal/mol
pre-miRNA408
>7H dna:primary_assembly
primary_assembly:MorexV3_pseudomolecules_assembly:7H:595182705:595182933:1
GAACAAGTGATAGAGATTGGATTTTGTGACTGGGGAGGGGAGAGGAGACAGGGATGGAGC
AGAGCAAGGGATGGGGCAAGCAACAAACTATCACCCCCTTATCATGAGAAGATCGAGAGA
GTTGTGAGAGACCAGGGATCCCTGTCGTCGTTGTTGTTCCTCCCTCCCTGCACTGCCTCT
TCCCTGGCTCCCCTCCCATCCCTCTCCATCTCCCTCTCTTGCTATTTTC

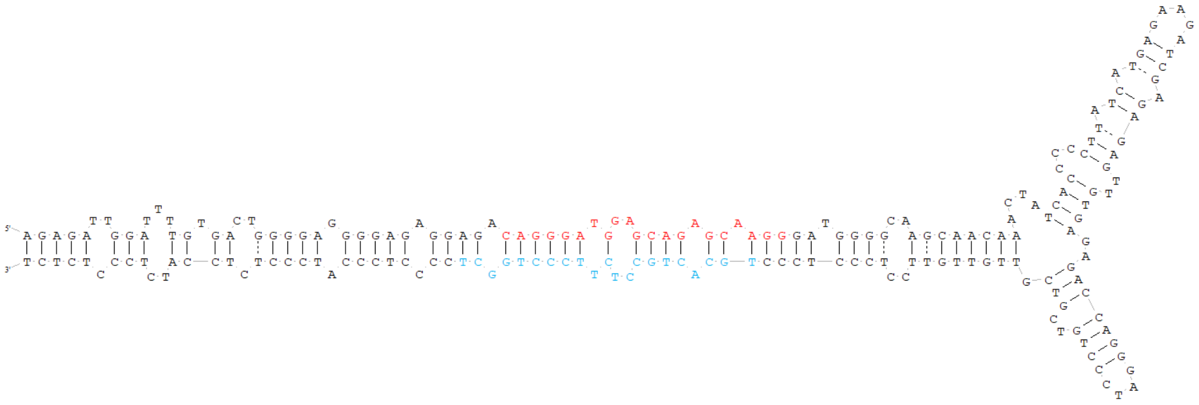

ΔG=-96.2 kcal/mol
pre-miRNA5052
>5H dna:primary_assembly
primary_assembly:MorexV3_pseudomolecules_assembly:5H:477138738:477139097:1
TGCCCAGAAGTATAAATTCACGGCTTTGATCTTTTTACTAACTTTGCCCTGATCTGACCC
TTGTTTATGATTTTTTTTTGGATATGATCCTTTTTTCTACCATAACCGGCTGGACGGTAG
GCATACACATCCTACCGCCACAAGCAGTAACTGTAGGGCTCGGATCAATGCGGATGGTTC
CTTCGGGTCAACGGAGTCAAACGACCTTATCGCCATTGTCTGCAGCGATAGGATGTGTAT
GCCTACCGCCATAGCGGGTGACGGTAGGAAAAAGTGTCAGATACAGAAAAAAAAATCGCA
AACCGAGTCGGGCCAAACTATGTAAAAAAAAGTCAAAACAGTGAAATTGCCCGCCTACGG

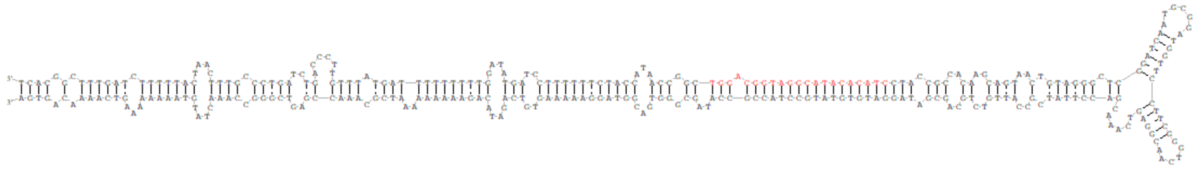

ΔG=-131.3 kcal/mol
pre-miRNA5168
>5H dna:primary_assembly
primary_assembly:MorexV3_pseudomolecules_assembly:5H:457064005:457064128:1
AAGTTAGGTTAAGGGGGTTGTTGTCTGGTTCAAGGTCTCCACATACATCATATACATGGA
GAGCATGGTATGGATTGTACGTTTTGGAGATCTCGGACCAGGCTTCAATCCCTTTAACCA
GCAT

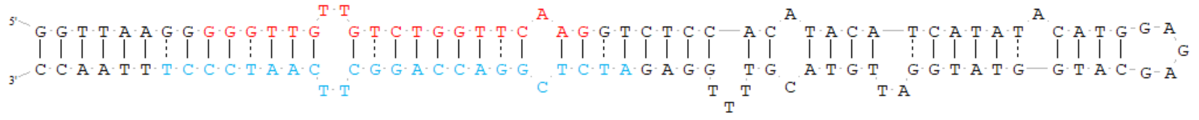

ΔG=-59.2 kcal/mol
pre-miRNA5200
>7H dna:primary_assembly
primary_assembly:MorexV3_pseudomolecules_assembly:7H:42107524:42107638:1
GTTTAAGCACCATTGCACGAAAGCCTTAGTGAATACCTACACTAGTAAGTTCTTCTTTAT
GTTGGAAGTGCTAGCGTTGTAGATACTCCCTAAGGCTTGGGTGGTATTGCGTGGT

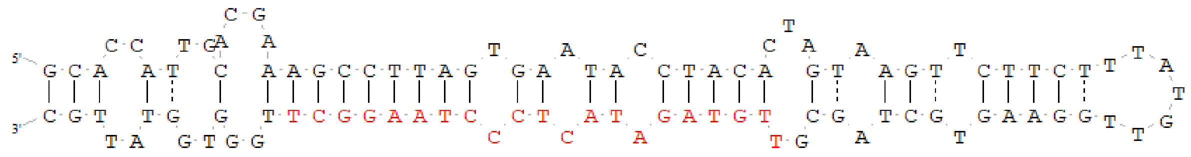

ΔG=-35.8 kcal/mol
pre-miRNA528
>6H dna:primary_assembly
primary_assembly:MorexV3_pseudomolecules_assembly:6H:92319425:92319801:1
GAGGCGAGGCCGAGGGGCGCGTGGGGCGGCCGGATCTGGCGGGGCGGCGAGGATGGAGGC
GCAGGCGCGCAGGGCGGCGAAGTGCGGCGGCGGCGGCACTTGGCAGTGCATGCGGCGAGG
CGGCGAAGTGCGGAACCGCCCGGCCGGGGCGGCGCGCTGCGGCGGCTGGGTGTACGTGGT

GTGCGGCTGACGGGAGGATTCCCCTATCCCTATCCTCGTTGGTGATTCCCCTATCCCTTC
CCTCTGTTTTTTCAATGGAACAGGGCTTAGCTTTGCTTTTTTGCTTGCCTCTCGGCTCTC
GCTCTCTCCTGTGCCTGCCTCTTCCATTCATGCCGCTAATGGCCGGCCGTGTCTCCATGC
GATGATGATTGGAGGTT

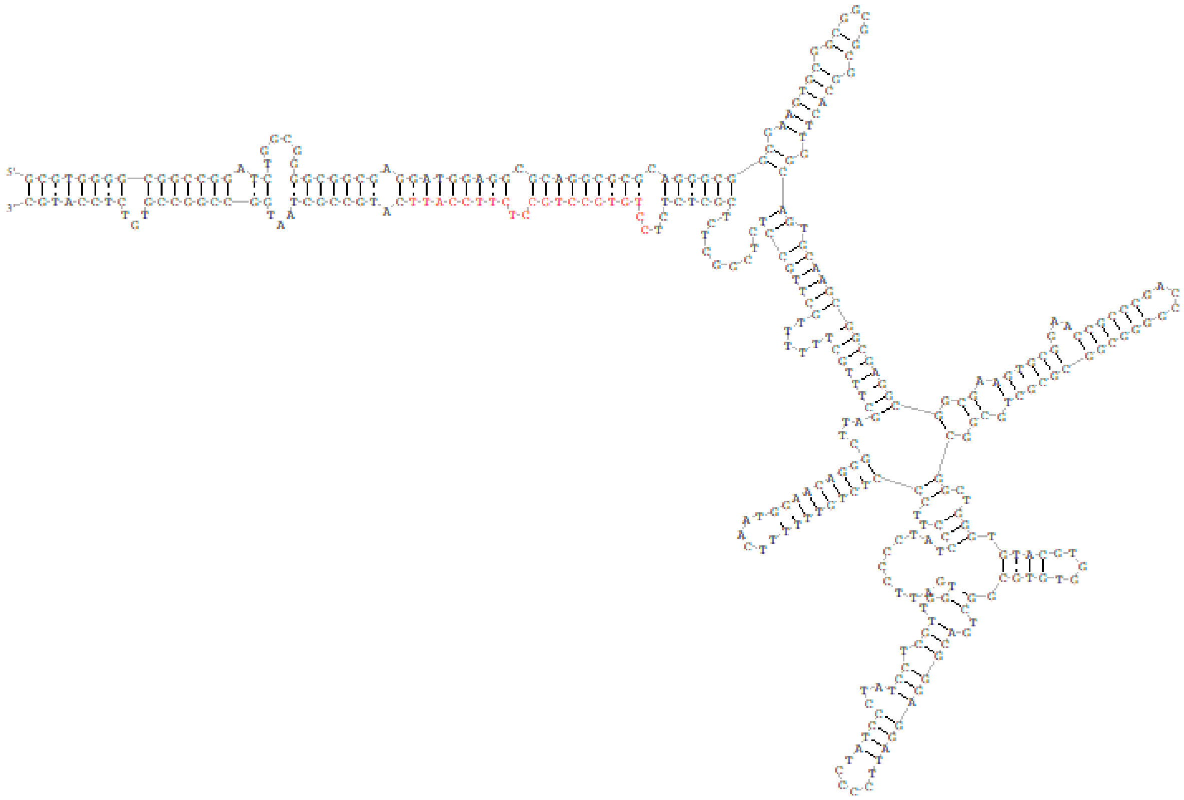

ΔG=-174.6 kcal/mol

pre-miRNA528

>4H dna:primary_assembly
primary_assembly:MorexV3_pseudomolecules_assembly:4H:589093145:589093275:-1
GCCGGAGCAGCAGCGGTGGAAGGGGCATGCAGAGGAGCGGCCATGCATGGGAGCTTTGCT
TTGCTTGCCTCTCCTGCTCTGGGCTCTAGCTCTCTCCTGTGCCTGCCTCTTCCATTCCTG
CCGCTAATGGC

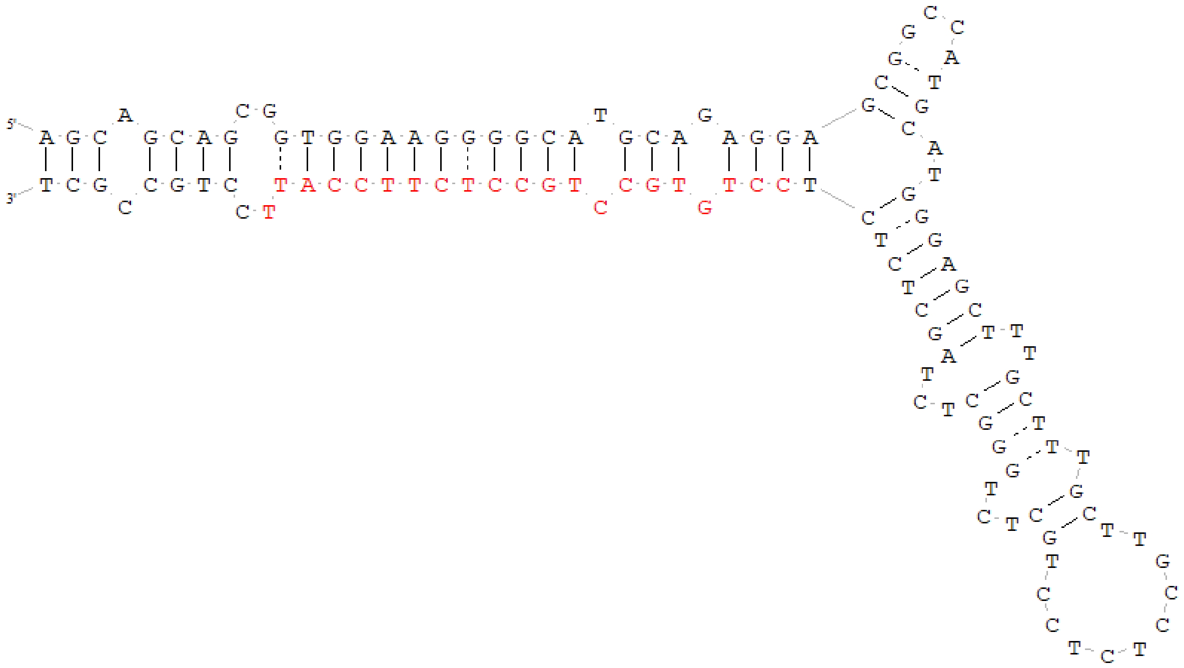

ΔG=-63.1 kcal/mol
pre-miRNA166c
>6H dna:primary_assembly
primary_assembly:MorexV3_pseudomolecules_assembly:6H:521815899:521816018:1
GAGCTTGTGGCCATGGCGGTTTGTGGGGAACGTTGGCTGGCTCGAGGCATCCTTGCCGAC
GGCGCGGGTTGAGGCGTCGGACCAGGCTTCATTCCTTTCAAACCGCCACGTCCATGCATC

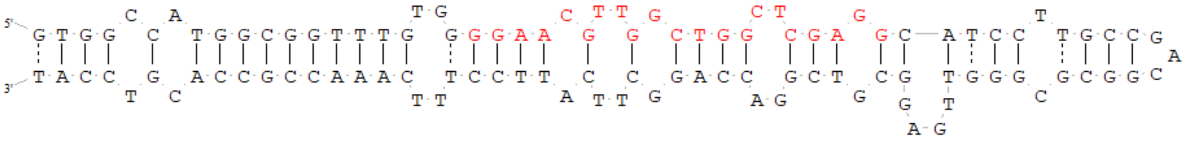

ΔG=-56 kcal/mol
pre-miRNA393
>2H dna:primary_assembly
primary_assembly:MorexV3_pseudomolecules_assembly:2H:655580455:655580551:1
TAGTGGAGGATTCCAAAGGGATCGCATTGATCGATCTCCCTCCGTCATCGGCGTGCAAGA
TCGATGGATCAGTGCGATCCCTCTGGAATTCTCCGCT

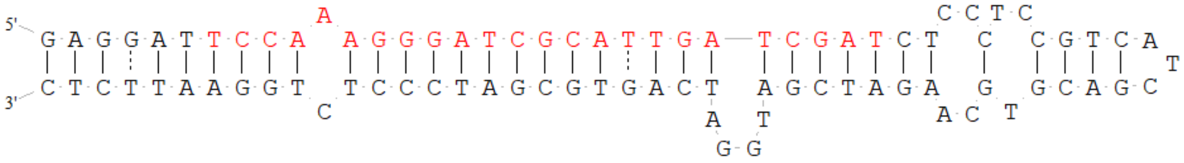

ΔG=-50.8 kcal/mol
pre-miRNA394_2
>6H dna:primary_assembly
primary_assembly:MorexV3_pseudomolecules_assembly:6H:426862934:426863191:1
CGTCCACGGAGGGGAGAGAGCTTGAAGTAAGGAAGTGGTAGTACTGGCCATATGGGCTTG
CCAAAGGGGCGCTTACCGAGAGCTCTTTGGCATTCTGTCCACCTCCTCCTGAATCTTGCA
GGGGAGATCTCACGATCTGTCGAGCTTGCTGATTGCTTGGAGGTGGGCATACTGCCAATG
GAGCTGTGTAGGCCTCCCTTTGTAAAACCCAGATGGCAAAAACCAACACGGGCTTCTTGC
TTAGTCTAGTTTTTCTCC

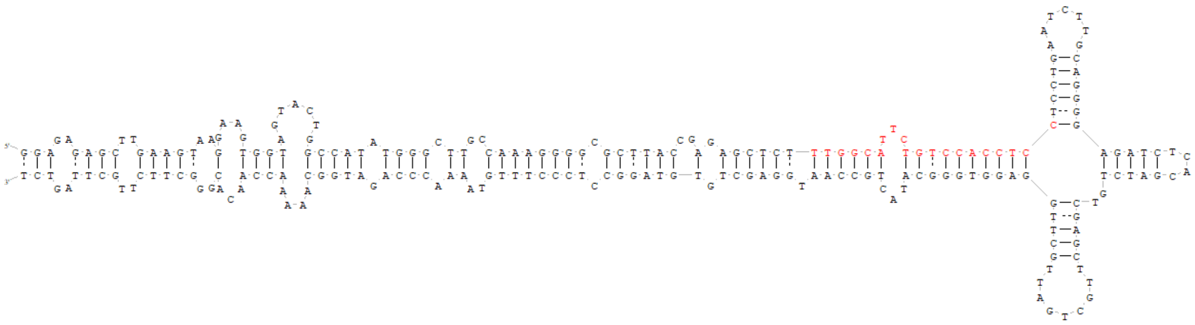

ΔG=-107.3 kcal/mol
pre-miRNA397a
>6H dna:primary_assembly
primary_assembly:MorexV3_pseudomolecules_assembly:6H:93800531:93800648:-1
GAGCCGCAGAGGTGCCGTTGAGTGCAGCGTTGATGAACCGTCCGGCCATGGCCCGTCCGC
CTCCACCGAGGCCGGAGCGGTTCACCGGCGCTGCACGCAATGACGCCTCTGCTTTCTC

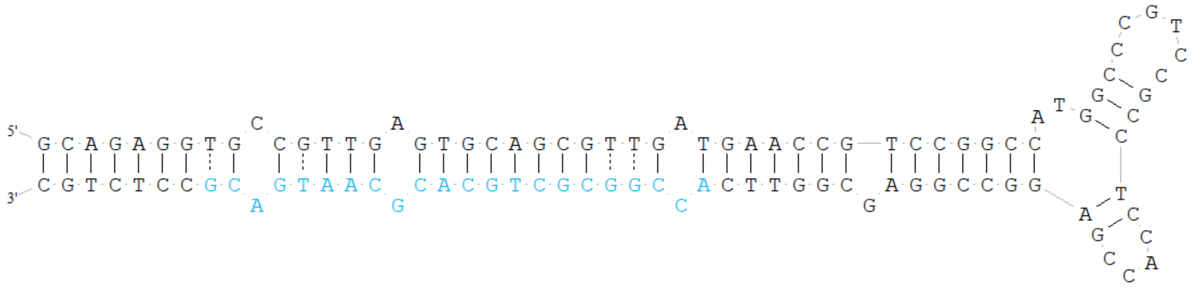

ΔG=-74.4 kcal/mol
pre-miRNA5051
>4H dna:primary_assembly
primary_assembly:MorexV3_pseudomolecules_assembly:4H:558076950:558077045:1
GTACTGTTCCAGTTTCAAGGTTTCAAAGCTGAAGGCTTCCCAGAGATCCTGCTGCTGGGT
GGTCTTTAGCTTTGGCACCTTGAAACTGGGACTTTA

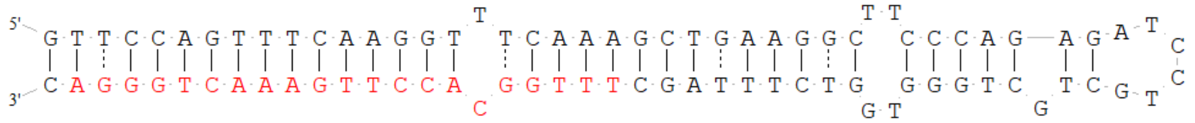

ΔG=-58.5 kcal/mol
pre-miRNA827
>2H dna:primary_assembly
primary_assembly:MorexV3_pseudomolecules_assembly:2H:544572786:544572977:-1
TCTGGTTCATGCAGCTCTGAACTTGTTTTGTTGGTTGTCATCTAACCATCATCGTTCCAT
CTGCCATCCGGTGCATGCATGGATGGACGCCTGAGTGAGTGCGTGGCGACTACGCCGACG
CACAGGCACAGACGGTGATGATGGTTAGATGACCATCAGCAAACATGTTCACGAGACATG
CGTCCTGGCAAC

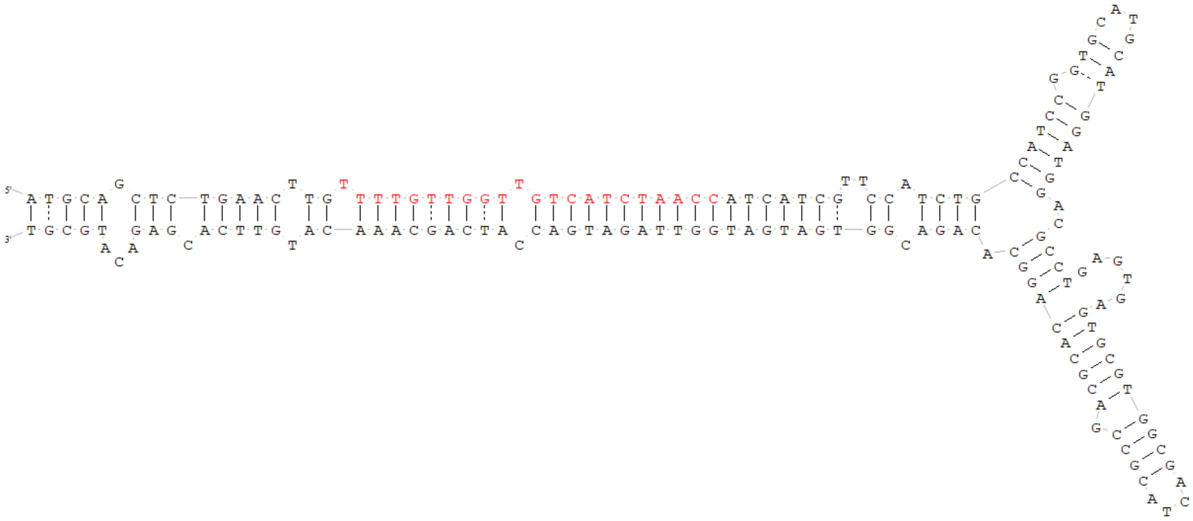

ΔG=-94.2 kcal/mol

pre-miRNA9662

>6H dna:primary_assembly
primary_assembly:MorexV3_pseudomolecules_assembly:6H:102023647:102023731:-1
CCCGGCGGCTCTGTGGTGTTCAAGCAGGAACCTCATGCTACCGGCAGCATGCGGCGCTTG
CTTGAACATCCCAGAGCCACCGGCG

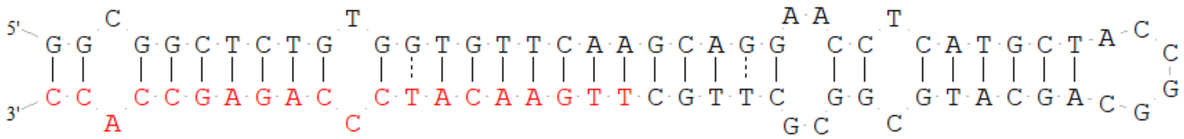

ΔG=-51.5 kcal/mol
