## Supplementary material for "Barley miRNAs and their targets regulation in response to heat stress at the early stage of development": Table S4

Table S4. Barley pri-miRNAs and cognate miRNAs with correlated changes in their levels under high temperature stress. Data marked in orange show upregulation of pri-miRNA/miRNA while data marked in blue show downregulation of pri-miRNA/miRNA levels. pri-miRNAs and miRNAs are grouped by miRNA families. *P* – pval * ≤0.05; ** ≤0.01; *** ≤0.001. FC, fold change.

| pri-miRNA [FC] | | | | | | | miRNA [FC] | | | |
|---|---|---|---|---|---|---|---|---|---|---|
| | 1h heat | *p* | 3h heat | *p* | 6h heat | *p* | | 1h heat | 3h heat | 6h heat |
| pri-miRNA159a | 1.67 | | 1.78 | ** | 2.32 | ** | miR159a-5p | 1.04 | 1.43 | 1.91 |
| | | | | | | | miR159a-3p | -1.95 | -6.27 | -5.75 |
| pri-miRNA160f | 1.09 | | 1.48 | ** | 1.38 | *** | miR160f | 1.58 | 1.55 | 1.62 |
| pri-miRNA166d | 1.74 | | 2.85 | *** | 3.71 | * | miR166d-5p | 1.44 | 2.34 | 1.21 |
| | | | | | | | miR166a/d-3p | 1.54 | 1.56 | 2.13 |
| pri-miRNA166a | 1.86 | * | 6.97 | * | 8.39 | * | miR166a-5p | -1.10 | -1.15 | 1.25 |
| pri-miRNA166e | 4.37 | ** | 7.20 | *** | 6.59 | ** | miR166a/d-3p | 1.54 | 1.56 | 2.13 |
| pri-miRNA166f | 2.65 | | 9.11 | | 6.84 | * | | | | |
| pri-miRNA166n | 1.36 | | 1.28 | | -1.50 | *** | miR166n-5p | -4.11 | -3.33 | -5.19 |
| pri-miRNA167c | 1.81 | | 1.36 | | 1.70 | | miR167b/c | 1.40 | -1.84 | 1.33 |
| pri-miRNA167b_1 | 1.43 | | 1.40 | | 2.63 | | | | | |
| pri-miRNA167d | 1.18 | | 1.20 | | 1.18 | | miR167b/d-5p | -1.09 | -1.21 | 1.08 |
| pri-miRNA167b_2 | 1.00 | | 1.11 | | 1.30 | | | | | |
| pri-miRNA169c | -2.05 | | -2.42 | *** | -2.90 | ** | miR169c.1-5p | -2.68 | -2.02 | -2.22 |
| | | | | | | | miR169c.2-5p | -1.03 | -1.67 | -1.19 |
| | | | | | | | miR169c-3p | -5.39 | -8.80 | -7.73 |
| pri-miRNA169d | -2.29 | * | -2.71 | * | -1.11 | | miR169d | -1.50 | -3.08 | -1.81 |
| pri-miRNA169i | -1.32 | | -2.53 | *** | -5.24 | *** | miR169e/g/i-5p | 1.12 | -1.50 | -1.15 |
| | | | | | | | miR169i-3p | -1.59 | -3.61 | -5.67 |
| pri-miRNA169e | 1.67 | | 1.31 | | -1.63 | | miR169e/g/i | 1.12 | -1.50 | -1.15 |
| pri-miRNA169i | 1.54 | | 1.58 | | -1.57 | | | | | |
| pri-miRNA169g | 1.90 | | 2.57 | | -1.46 | | | | | |
| pri-miRNA169h | 1.03 | | -1.43 | *** | -3.04 | ** | miR169h-5p | -1.19 | -1.02 | -2.53 |
| | | | | | | | miR169h-3p | 1.00 | -1.74 | -4.28 |
| pri-miRNA172b | 2.36 | | 2.89 | *** | 5.06 | *** | miR172b-5p | -1.83 | 1.00 | 2.22 |
| pri-miRNA319b | 1.65 | | 1.55 | ** | 2.51 | *** | miR319b.1-5p | 2.95 | 1.34 | 11.59 |
| | | | | | | | miR319b.2-5p | -4.80 | -11.46 | -3.27 |
| | | | | | | | miR319b-3p | 1.21 | 1.38 | 1.69 |
| pri-miRNA408 | 1.26 | | -3.78 | *** | -1.99 | ** | miR408-5p | -1.78 | -9.57 | -5.44 |
| | | | | | | | miR408-3p | 1.08 | -1.93 | 1.05 |
| pri-miRNA444a | -1.81 | ** | 2.32 | | 2.11 | * | miR444a.2 | 2.65 | 2.86 | 2.41 |
| pri-miRNA5052 | -3.05 | * | -3.58 | ** | -4.72 | *** | miR5052 | -1.80 | -1.32 | -4.66 |
| pri-miRNA5168 | 2.29 | * | 3.49 | ** | 3.05 | * | miR5168-5p | 1.05 | 1.98 | 2.04 |
| | | | | | | | miR5168-3p | 1.01 | -1.06 | -1.10 |
| pri-miRNA5200 | -10.76 | *** | -7.12 | *** | -4.47 | *** | miR5200 | -3.90 | -5.37 | -19.06 |
| pri-miRNA528 | -1.78 | | -9.85 | *** | -6.56 | *** | miR528 | -2.75 | -7.07 | -6.14 |
| pri-miRNA528 | -1.80 | | -10.71 | *** | -2.82 | ** | | | | |
| pri-miRNAx9a | 1.01 | | 1.28 | | -1.07 | | miRx9a | 1.28 | -1.54 | -1.51 |
