## Supplementary material for "Barley miRNAs and their targets regulation in response to heat stress at the early stage of development": Table S5

Table S5. Barley pri-miRNAs and cognate miRNAs that show no correlation in their level under high temperature stress. Data marked in orange show upregulation of pri-miRNA/miRNA while data marked in blue show downregulation of pri-miRNA/miRNA levels. pri-miRNAs and miRNAs are grouped by miRNA families. *P* – pval * ≤0.05; **≤ 0.01; *** ≤0.001. FC, fold change.

| | pri-miRNA [FC] | | | | | | | miRNA [FC] | | |
|---|---|---|---|---|---|---|---|---|---|---|
| | 1h heat | *p* | 3h heat | *p* | 6h heat | *p* | | 1h heat | 3h heat | 6h heat |
| pri-miRNA1130 | 2.55 | | 2.81 | | 3.05 | *** | miR1130 | 2.59 | 1.46 | -1.49 |
| pri-miRNA1432 | 1.59 | | -2.06 | * | -2.40 | * | miR1432-5p | 1.18 | -1.65 | 1.18 |
| pri-miRNA159b | 1.36 | | 2.30 | | 4.39 | ** | miR159b | 1.26 | 2.26 | -1.25 |
| pri-miRNA159a | 1.43 | | 2.83 | ** | 3.58 | *** | | | | |
| pri-miRNA160c_1 | 2.15 | | 2.50 | * | 3.28 | * | miR160a | -1.02 | -2.00 | -1.31 |
| pri-miRNA160b | 2.09 | | 2.56 | | 2.36 | * | | | | |
| pri-miRNA160c_2 | 3.57 | | 2.70 | * | 3.22 | | | | | |
| pri-miRNA160a | 13.65 | | 2.66 | ** | 4.03 | * | | | | |
| pri-miRNA160e | -1.20 | | 1.29 | | 1.18 | | miRNA160e | -1.03 | -2.50 | -1.65 |
| pri-miRNA166c | 1.14 | | 1.29 | | 1.70 | | miR166c-5p | -1.96 | -3.90 | -2.80 |
| | | | | | | | miR166c-3p | 1.25 | -1.18 | -1.22 |
| pri-miRNA169a | 1.92 | | 1.49 | | 3.69 | * | miR169a | 2.43 | -2.56 | 1.60 |
| pri-miRNA171a | 6.73 | | 14.78 | | 4.34 | | miR171a/b/e | 1.24 | -1.46 | 1.15 |
| pri-miRNA171b | 1.61 | | 1.63 | | 1.23 | | | | | |
| pri-miRNA171e | 2.61 | | 5.20 | * | 5.45 | ** | | | | |
| pri-miRNA171c | 3.41 | * | 4.78 | ** | 3.89 | ** | miR171c | 1.57 | -1.32 | -1.37 |
| pri-miRNA171d | 1.58 | | 4.25 | ** | 7.07 | ** | miR171d | 1.17 | -1.05 | -1.15 |
| pri-miRNA171h | -1.19 | | 2.45 | | -1.38 | | miR171h | -1.39 | -1.90 | -1.57 |
| pri-miRNA393 | 2.23 | | 3.17 | *** | 2.90 | ** | miR393 | -1.02 | -1.43 | 1.02 |
| pri-miRNA394_2 | 2.99 | * | 5.22 | | 3.22 | * | miR394 | 1.22 | -1.31 | -1.54 |
| pri-miRNA394_1 | 2.16 | | 5.00 | | 3.62 | | | | | |
| pri-miRNA395f | 2.42 | | 6.19 | * | 41.64 | * | miR395b/f | 1.31 | 2.23 | 1.73 |
| pri-miRNA395b | 6.74 | | 41.38 | | 32.68 | * | | | | |
| pri-miRNA396a | 2.26 | * | 3.17 | ** | 3.44 | ** | miR396a | -1.11 | 1.63 | 1.34 |
| pri-miRNA396c | 2.33 | | -1.02 | | 2.22 | * | miR396c-5p | 1.04 | 1.26 | -1.11 |
| pri-miRNA396e | 2.46 | | 5.08 | * | 4.32 | | miR396e-5p | -1.20 | 1.35 | 1.35 |
| pri-miRNA396h | 1.40 | | -1.35 | | 1.84 | | miR396h | -1.29 | 2.15 | 1.98 |
| pri-miRNA397a | -1.02 | | -1.21 | | 1.08 | | miR397a-3p | -1.45 | -8.67 | -4.57 |
| pri-miRNA398f | 18.02 | | 2.36 | | 2.20 | *** | miR398f | 1.19 | -1.33 | 1.02 |
| pri-miRNA399g_1 | 1.31 | | -2.00 | *** | -2.43 | ** | miR399d/g | -1.14 | -1.34 | -1.86 |
| pri-miRNA399d_1 | -1.02 | | -5.20 | ** | 2.81 | | | | | |
| pri-miRNA399g_2 | 1.79 | | -3.61 | ** | -2.09 | | | | | |
| pri-miRNA399d_2 | 1.83 | | -1.18 | | 1.00 | | | | | |
| pri-miRNA399h | 1.13 | | 2.32 | | 2.23 | | miR399h | 1.40 | -1.63 | -1.81 |
| pri-miRNA444b | -1.93 | * | -1.81 | *** | -1.30 | | miR444b | -1.21 | 1.18 | -1.21 |
| pri-miRNA444c | 10.40 | | 1.37 | | 1.38 | * | miR444c | -1.45 | -1.79 | -1.43 |
| pri-miRNA5048b | 1.00 | | 1.28 | * | 1.17 | | miR5048b | 1.09 | 1.23 | -1.13 |
| pri-miRNA5049b | -1.27 | | -1.51 | *** | 1.12 | | miR5049b | 1.26 | -1.65 | -2.36 |

| | | | | | | | | | |
|---|---|---|---|---|---|---|---|---|---|
| pri-miRNA5049e | 1.56 | 1.76 | * | 1.69 | | miR5049e | -1.48 | 1.16 | 1.42 |
| pri-miRNA5051 | -1.67 | -1.36 | * | -1.12 | | miR5051 | -2.85 | 1.29 | 2.10 |
| pri-miRNA827 | 1.13 | -1.18 | | -1.97 | | miR827-5p | -1.59 | -2.82 | -3.15 |
| pri-miRNA9662 | -1.18 | 1.45 | | 1.31 | | miR9662 | 1.25 | 2.00 | 2.24 |
| pri-miRNAx9b | 1.08 | 1.68 | *** | 1.62 | ** | miRx9b | 1.12 | -1.41 | -1.35 |
| pri-miRNAx13 | 1.53 | 4.46 | * | 4.85 | * | miRx13 | 1.01 | -1.49 | -3.76 |
