## Supplementary material for "Barley miRNAs and their targets regulation in response to heat stress at the early stage of development": Table S6

Table S6. List of barley target mRNAs identified for miRNAs most prominently changed during heat stress treatment using degradome data.

| miRNA | miR sequence | miRNA length | Gene ID | Description |
|---|---|---|---|---|
| miR156_1 | TGACAGAAGAGAGTGAGCACA | 21 | HORVU0Hr1G039170 | Squamosa promoter-binding-like protein 16 |
| | | | HORVU3Hr1G094730 | Squamosa promoter-binding-like protein 2 |
| | | | HORVU6Hr1G031450 | squamosa promoter binding protein-like 2 |
| miR159_1 | CTTGGACTGAAGGGTGCTCC | 20 | HORVU3Hr1G079490 | myb domain protein 33 |
| | | | HORVU7Hr1G090120 | Transcription factor GAMYB |
| | | | HORVU3Hr1G079490 | myb domain protein 33 |
| miR160a | TGCCTGGCTCCCTGTATGCCA | 21 | HORVU1Hr1G041770 | Auxin response factor 22 |
| | | | HORVU2Hr1G089670 | auxin response factor 10 |
| | | | HORVU6Hr1G026750 | Auxin response factor 18 |
| | | | HORVU6Hr1G058890 | auxin response factor 16 |
| | | | HORVU7Hr1G101270 | auxin response factor 16 |
| miR160e | TGCCTGGCTCCCTGAATGCCA | 21 | HORVU1Hr1G041770 | Auxin response factor 22 |
| | | | HORVU2Hr1G089670 | auxin response factor 10 |
| | | | HORVU6Hr1G026750 | Auxin response factor 18 |
| | | | HORVU6Hr1G058890 | auxin response factor 16 |
| | | | HORVU7Hr1G101270 | auxin response factor 16 |
| miR160_1 | GTGCCTGGCTCCCTGTATGCC | 21 | HORVU6Hr1G026750 | Auxin response factor 18 |
| | | | HORVU7Hr1G101270 | auxin response factor 16 |
| miR165a/b | TCGGACCAGGCTTCATCCCCC | 21 | HORVU0Hr1G010250 | Homeobox-leucine zipper protein HOX32 |
| | | | HORVU1Hr1G041790 | Homeobox-leucine zipper protein family |
| | | | HORVU3Hr1G026990 | Homeobox-leucine zipper protein family |
| | | | HORVU5Hr1G010650 | Homeobox-leucine zipper protein family |
| | | | HORVU5Hr1G093150 | Homeobox-leucine zipper protein HOX32 |
| miR166a | TCGGACCAGGCTTCATTCCCCT | 21 | HORVU0Hr1G010250 | Homeobox-leucine zipper protein HOX32 |
| | | | HORVU1Hr1G041790 | Homeobox-leucine zipper protein family |
| | | | HORVU2Hr1G045590 | unknown function |
| | | | HORVU3Hr1G026990 | Homeobox-leucine zipper protein family |
| | | | HORVU5Hr1G010650 | Homeobox-leucine zipper protein family |
| | | | HORVU5Hr1G093150 | Homeobox-leucine zipper protein HOX32 |
| miR166a/d-3p | CTCGGACCAGGCTTCATTCC | 20 | HORVU0Hr1G010250 | Homeobox-leucine zipper protein HOX32 |
| | | | HORVU3Hr1G026990 | Homeobox-leucine zipper protein family |
| | | | HORVU5Hr1G010650 | Homeobox-leucine zipper protein family |
| miR166c-3p | TCGGACCAGGCTTCATTCCTT | 21 | HORVU0Hr1G010250 | Homeobox-leucine zipper protein HOX32 |
| | | | HORVU1Hr1G041790 | Homeobox-leucine zipper protein family |

| | | | | |
|---|---|---|---|---|
| | | | HORVU2Hr1G045590 | unknown function |
| | | | HORVU3Hr1G026990 | Homeobox-leucine zipper protein family |
| | | | HORVU5Hr1G010650 | Homeobox-leucine zipper protein family |
| | | | HORVU5Hr1G093150 | Homeobox-leucine zipper protein HOX32 |
| miR166g | TTCGGACCAGGCTTCAATCCC | 21 | HORVU0Hr1G010250 | Homeobox-leucine zipper protein HOX32 |
| | | | HORVU3Hr1G026990 | Homeobox-leucine zipper protein HOX32 |
| | | | HORVU5Hr1G010650 | Homeobox-leucine zipper protein family |
| miR166_2 | TCGGACCAGGCTTCAATCCCT | 21 | HORVU1Hr1G041790 | Homeobox-leucine zipper protein family |
| | | | HORVU3Hr1G026990 | Homeobox-leucine zipper protein family |
| | | | HORVU5Hr1G010650 | Homeobox-leucine zipper protein family |
| | | | HORVU5Hr1G093150 | Homeobox-leucine zipper protein HOX32 |
| | | | HORVU0Hr1G010250 | Homeobox-leucine zipper protein HOX32 |
| miR167b/c | TGAAGCTGCCAGCATGATCTA | 21 | HORVU2Hr1G121110 | auxin response factor 6 |
| | | | HORVU5Hr1G009650 | auxin response factor 6 |
| | | | HORVU6Hr1G026730 | auxin response factor 6 |
| | | | HORVU7Hr1G106280 | auxin response factor 6 |
| miR167b/d-5p | TGAAGCTGCCAGCATGATCTG | 21 | HORVU2Hr1G121110 | auxin response factor 6 |
| | | | HORVU5Hr1G009650 | auxin response factor 6 |
| | | | HORVU6Hr1G026730 | auxin response factor 6 |
| | | | HORVU7Hr1G106280 | auxin response factor 6 |
| miR167b/d-5p | TGAAGCTGCCAGCATGATCTGA | 22 | HORVU2Hr1G121110 | auxin response factor 6 |
| | | | HORVU5Hr1G009650 | auxin response factor 6 |
| | | | HORVU7Hr1G106280 | auxin response factor 6 |
| miR168 | TCGCTTGGTGCAGATCGGGACC | 22 | HORVU7Hr1G007000 | Argonaute family protein |
| miR169a | CAGCCAAGGATGACTTGCCGA | 21 | HORVU2Hr1G032130 | Nuclear transcription factor Y subunit A-5 |
| | | | HORVU4Hr1G075830 | Nuclear transcription factor Y subunit A-3 |
| | | | HORVU6Hr1G081080 | Nuclear transcription factor Y subunit A-5 |
| miR169c.2-5p | CAGCCAAGGATGACTTGCCGG | 21 | HORVU2Hr1G032130 | Nuclear transcription factor Y subunit A-5 |
| | | | HORVU4Hr1G075830 | Nuclear transcription factor Y subunit A-3 |
| | | | HORVU6Hr1G081080 | Nuclear transcription factor Y subunit A-5 |
| miR169d | TAGCCAAGAATGACTTGCCT | 20 | HORVU4Hr1G005670 | Nuclear transcription factor Y subunit A-9 |
| | | | HORVU5Hr1G007890 | Nuclear transcription factor Y subunit A-10 |
| miR169e/g/i-5p | TAGCCAAGGATGACTTGCCTG | 21 | HORVU2Hr1G032130 | Nuclear transcription factor Y subunit A-5 |
| | | | HORVU4Hr1G005670 | Nuclear transcription factor Y subunit A-9 |
| | | | HORVU4Hr1G075830 | Nuclear transcription factor Y subunit A-3 |
| | | | HORVU5Hr1G007890 | Nuclear transcription factor Y subunit A-10 |

| | | | | |
|---|---|---|---|---|
| | | | HORVU6Hr1G081080 | Nuclear transcription factor Y subunit A-5 |
| miR169h-5p | TAGCCAAGGATGACTTGCCTA | 21 | HORVU2Hr1G032130 | Nuclear transcription factor Y subunit A-5 |
| | | | HORVU4Hr1G075830 | Nuclear transcription factor Y subunit A-3 |
| | | | HORVU5Hr1G007890 | Nuclear transcription factor Y subunit A-10 |
| | | | HORVU6Hr1G081080 | Nuclear transcription factor Y subunit A-5 |
| miR171a/b/e | TGATTGAGCCGTGCCAATATC | 21 | HORVU6Hr1G063650 | GRAS family transcription factor |
| miR171c | TTGAGCCGTGCCAATATCACG | 21 | HORVU6Hr1G063650 | GRAS family transcription factor |
| miR171d | TGATTGAGCCGCGCCAATATC | 21 | HORVU6Hr1G063650 | GRAS family transcription factor |
| miR319b.1-5p | TTGGACTGAAGGGTGCTCCCT | 21 | HORVU2Hr1G060120 | TCP family transcription factor 4 |
| | | | HORVU3Hr1G028240 | Transcription factor TCP4 |
| | | | HORVU5Hr1G103400 | TCP family transcription factor 4 |
| miR319b-3p | CTTGGACTGAAGGGTGCTCCCT | 22 | HORVU2Hr1G060120 | TCP family transcription factor 4 |
| miR319b | CTTGGACTGAAGGGTGCTCC | 20 | HORVU2Hr1G060120 | TCP family transcription factor 4 |
| miR319_2 | TTGGACTGAAGGGAGCTCCCT | 21 | HORVU2Hr1G060120 | TCP family transcription factor 4 |
| | | | HORVU3Hr1G028240 | Transcription factor TCP4 |
| | | | HORVU5Hr1G103400 | TCP family transcription factor 4 |
| miR390 | CGCTATCTATCCTGAGCTCC | 20 | HORVU1Hr1G068940 | Ribosomal protein L34e superfamily protein |
| miR393_1 | TCCAAAGGGATCGCATTGATC | 21 | HORVU1Hr1G021550 | Transport inhibitor response 1-like protein |
| miR393_2 | TTCCAAAGGGATCGCATTGAT | 21 | HORVU1Hr1G021550 | Transport inhibitor response 1-like protein |
| | | | HORVU2Hr1G070800 | Transport inhibitor response 1-like protein |
| | | | HORVU6Hr1G038740 | CLP1-similar protein 5 |
| miR396a | TCCACAGGCTTTCTTGAACTG | 21 | HORVU0Hr1G016590 | growth-regulating factor 3 |
| | | | HORVU0Hr1G016610 | Growth-regulating factor 4 |
| | | | HORVU0Hr1G026650 | Growth-regulating factor 4 |
| | | | HORVU2Hr1G101770 | Growth-regulating factor 3 |
| | | | HORVU4Hr1G003440 | Growth-regulating factor 9 |
| | | | HORVU6Hr1G068370 | Growth-regulating factor 4 |
| | | | HORVU7Hr1G008680 | growth-regulating factor 5 |
| miR396c-5p | TTCCACAGCTTTCTTGAACTT | 21 | HORVU2Hr1G101770 | Growth-regulating factor 3 |
| | | | HORVU4Hr1G003440 | Growth-regulating factor 9 |
| | | | HORVU4Hr1G037480 | Growth-regulating factor 8 |
| miR396e-5p | TTCCACAGCTTTCTTGAACTG | 21 | HORVU0Hr1G016590 | growth-regulating factor 3 |
| | | | HORVU0Hr1G026650 | Growth-regulating factor 4 |
| | | | HORVU2Hr1G101770 | Growth-regulating factor 3 |
| | | | HORVU4Hr1G003440 | Growth-regulating factor 9 |
| | | | HORVU6Hr1G068370 | Growth-regulating factor 4 |
| | | | HORVU7Hr1G008680 | growth-regulating factor 5 |
| miR396h | TCCACAGGCTTTCTTGAACGG | 21 | HORVU0Hr1G016590 | growth-regulating factor 3 |
| | | | HORVU0Hr1G016610 | Growth-regulating factor 4 |

| | | | | |
|---|---|---|---|---|
| | | | HORVU0Hr1G026650 | Growth-regulating factor 4 |
| | | | HORVU2Hr1G101770 | Growth-regulating factor 3 |
| | | | HORVU4Hr1G003440 | Growth-regulating factor 9 |
| | | | HORVU6Hr1G068370 | Growth-regulating factor 4 |
| | | | HORVU7Hr1G008680 | growth-regulating factor 5 |
| miR399d/g | TGCCAAAGGAGATTTGCCCCG | 21 | HORVU1Hr1G085570 | ubiquitin-conjugating enzyme 23 |
| miR399h | TGCCAAAGGAGAATTGCCCTG | 21 | HORVU1Hr1G085570 | ubiquitin-conjugating enzyme 23 |
| miR444a | TTGTGGCTTTCTTGCAAGTC | 20 | HORVU6Hr1G073040 | MADS-box transcription factor 57 |
| miR444a | TGCAGTTGCTGCCTCAAGCT | 20 | HORVU6Hr1G073040 | MADS-box transcription factor 57 |
| | | | HORVU7Hr1G066380 | zinc finger (C3HC4-type RING finger) family protein |
| miR444a.2 | TGCAGTTGCTGCCTCAAGCTT | 21 | HORVU6Hr1G073040 | MADS-box transcription factor 57 |
| | | | HORVU7Hr1G066380 | zinc finger (C3HC4-type RING finger) family protein |
| miR444b | TGCAGTTGCTGTCTCAAGCTT | 21 | HORVU6Hr1G073040 | MADS-box transcription factor 57 |
| | | | HORVU7Hr1G066380 | zinc finger (C3HC4-type RING finger) family protein |
| miR827-5p | TTAGATGACCATCAGCAAACA | 21 | HORVU2Hr1G094690 | SPX domain-containing membrane protein |
| miR1122a | AGTCTTTTTAGAGATTCCACT | 21 | HORVU1Hr1G060100 | SAGA-associated factor 11 (histone acetyltransferase complex) |
| | | | HORVU3Hr1G079210 | calmodulin 5 |
| miR1432-5p | TTCAGGAGAGATGACACCGAC | 21 | HORVU0Hr1G005300 | 2-oxoglutarate (2OG) and Fe(II)-dependent oxygenase superfamily protein |
| | | | HORVU1Hr1G094160 | calmodulin like 43 |
| | | | HORVU3Hr1G016490 | tryptophan aminotransferase related 2 |
| miR1436_1 | ATTATGGGACGGAGGGAGTAG | 21 | HORVU1Hr1G054460 | Phosphatidylinositol N-acetylglucosaminyltransfera se subunit P |
| | | | HORVU2Hr1G026190 | Benzoate O-methyltransferase |
| | | | HORVU5Hr1G123680 | RNA-binding protein 34 |
| | | | HORVU7Hr1G122690 | Phosphatidylinositol-4-phosphate 5-kinase family protein |
| miR5048b | TATTTGCAGGTTTTAGGTCTAA | 22 | HORVU7Hr1G001600 | cysteine-rich RLK (RECEPTOR-like protein kinase) 33 |
| miR5049b | AGTATTTAGGTACAGAGGGAG | 21 | HORVU2Hr1G039520 | Cytochrome P450 superfamily protein |
| | | | HORVU4Hr1G025070 | Protein kinase superfamily protein |
| miR5049_5 | AATTAATATGGATCGGAGGGA | 21 | HORVU6Hr1G009380 | Protein canopy-1 |
| miR5175a | AAGAATTTTGGGACGGAGGGA | 21 | HORVU7Hr1G097920 | Pentatricopeptide repeat-containing protein |
| miR6191 | TGTCTTAGATTTGTCTAGATA | 21 | HORVU3Hr1G096140 | Exostosin family protein |
| | | | HORVU5Hr1G018050 | kinesin 5 |
| miR6197_1 | TCTGTTCCTAAATGTAAGACG | 21 | HORVU4Hr1G084110 | Cysteine-rich repeat secretory protein 11 |
| | | | HORVU7Hr1G103180 | D.melanogaster polytene |

| | | | | |
|---|---|---|---|---|
| miR9662 | TTGAACATCCCAGAGCCACC | 20 | HORVU5Hr1G109600 | Mitochondrial transcription termination factor family protein |
| | | | HORVU6Hr1G003560 | Mitochondrial transcription termination factor family protein |
| | | | HORVU6Hr1G005650 | Mitochondrial transcription termination factor family protein |
| | | | HORVU7Hr1G037730 | Mitochondrial transcription termination factor family protein |
| | | | HORVU7Hr1G040960 | Mitochondrial transcription termination factor family protein |
| miR9863a/b | TGAGAAGGTAGATCATAATAGC | 22 | HORVU3Hr1G105020 | Disease resistance protein |
